# Circular RNA vaccine performance is determined by RNA quality and epitranscriptomic tuning rather than innate activation

**DOI:** 10.64898/2026.06.11.731499

**Authors:** Abby L Martin, Jessica F Cotterell, Kaitlin H Buick, Thomas W Bird, Joanna Kuang, Olga R Palmer, Ngarangi C Mason, Lydia White, Sarah L Draper, Amy J Foster, Gavin F Painter, Isabelle Montgomerie, Tifany Oulavallickal, Wayne M Patrick, Lisa M Connor

**Affiliations:** Malaghan Institute of Medical Research, Wellington, New Zealand; School of Biological Sciences, Victoria University of Wellington, Wellington New Zealand; Ferrier Research Institute, Victoria University of Wellington, Wellington, New Zealand

## Abstract

Circular RNA (circRNA) is an emerging vaccine modality that is proposed to improve stability, reduce reactogenicity and extend antigen expression compared with linear mRNA. However, the relative contributions to vaccine performance of its covalently closed structure, its purity, and of nucleotide modifications remain poorly described. Here, we systematically dissected these parameters in vivo across two antigen systems. We identified RNA quality as a major determinant of circRNA reactogenicity, with differences in innate immune activation tracking with the presence of residual RNA species in less refined preparations. In contrast, highly purified circRNA exhibited markedly reduced reactogenicity compared with linear mRNA, independent of nucleotide modification. Despite these differences, circRNA and mRNA vaccines elicited comparable antibody titres and T cell responses, indicating that reduced innate activation does not enhance adaptive immune magnitude. Notably, incorporating *N*^6^-methyladenosine (m^6^A) did not affect reactogenicity or antigen expression but selectively enhanced antibody quality, increasing binding affinity and neutralisation capacity. CircRNA vaccination also altered the anatomical distribution of germinal centre responses, reducing splenic antigen-specific germinal centre B cells while preserving lymph node responses. Together, these findings show that circRNA vaccine performance is governed by RNA preparation quality and epitranscriptomic tuning rather than innate activation alone.

## Introduction

Circular RNA (circRNA) is characterised by a covalently closed structure, which confers resistance to exonuclease-mediated degradation and enhances RNA stability^1–3^. Unlike linear RNA, circRNA lacks free 5′ and 3′ termini, which are canonical recognition features for cytosolic RNA sensors such as RIG-I^4–7^, although innate sensing of circRNA remains context dependent and can vary with RNA sequence, structure and purity^8^. These properties have led to the proposal that circRNA vaccines may exhibit reduced reactogenicity and support more durable antigen expression than linear mRNA platforms^1,2,9^. However, the relative contributions of its covalently closed architecture, manufacturing purity, and nucleotide modification to circRNA vaccine performance remain incompletely defined.

After demonstrating rapid scalability and high efficacy against SARS-CoV-2^10–12^, mRNA vaccines have continued to transform the vaccinology landscape, across both infectious diseases^13,14^ and cancer^15,16^. These advances have highlighted the importance of RNA chemistry and manufacturing in shaping both reactogenicity (innate immune initiation) and immunogenicity (vaccine-associated immune responses). Early studies of mRNA transcribed in vitro showed that double-stranded RNA contaminants generated during transcription strongly activate RIG-I and MDA5, triggering innate antiviral signalling pathways, such as the type I interferons that suppress mRNA translation and reduce protein production^17–19^. A reduction in protein (antigen) expression can decrease vaccine potency while simultaneously contributing to inflammatory reactogenicity. To mitigate these effects, incorporating modified nucleotides, including *N*^1^-methylpseudouridine (m1Ψ)^19,20^, together with high-performance liquid chromatography (HPLC) purification^21^, have become central to optimising clinical mRNA vaccine platforms.

In contrast to linear mRNA, circRNA production requires a circularisation step, most commonly achieved using permuted intron-exon (PIE) strategies in which a linear precursor undergoes autocatalytic splicing to generate a covalently closed RNA circle^3,22^. This process can generate heterogenous RNA populations, including non-circularised precursor RNA, excised intron fragments, nicked circles, and other products that need to be removed during downstream purification. These species can influence both innate immune sensing and translational efficiency. Furthermore, circRNA lacks a 5′ cap and poly(A) tail, instead relying on cap-independent translation, typically driven by an internal ribosome entry site (IRES)^3,23^. As a result, circRNA vaccine performance is not only shaped by its covalently closed topology, but also by design features and RNA quality, which together influence antigen expression and immunological outcomes.

Circular RNA vaccines have shown promising efficacy across multiple infectious disease models^9^. Studies with SARS-CoV-2 have demonstrated robust neutralising antibody and T cell responses in mice and non-human primates^2,9,24,25^, with similar findings reported for Zika virus, influenza, and other viral pathogens^26–29^. However, whether circRNA is intrinsically less reactogenic than linear mRNA remains unresolved. While some reports have shown that purified circRNA exhibits reduced innate immune activation and prolonged translation^1,3^, other studies have shown that exogenous circRNA can still activate RIG-I-dependent sensing pathways^8^, indicating that a circular structure alone does not abolish immune recognition. Emerging evidence suggests that reactogenicity is influenced by multiple factors, including RNA sequence and structure^30^, the circularisation strategy^31,32^, and the presence of impurities^33^. In addition, *N*^6^-methyladenosine (m^6^A) has been implicated in both circRNA translation and immune recognition, with studies suggesting that m^6^A modification can dampen RIG-I-mediated innate activation while promoting cap-independent translation of circRNA^34,35^.

Here, we systematically dissect these factors using a controlled in vivo framework to directly compare circRNA and linear mRNA vaccines. Our findings reveal that circRNA reactogenicity is primarily driven by the quality of its preparation, rather than circular topology alone, and that, when highly purified, circRNA exhibits reduced innate immune activation and prolonged antigen expression compared with linear mRNA without enhancing overall immune magnitude. Instead, circRNA vaccination promotes distinct lymphoid tissue-specific B cell responses, while m^6^A modification selectively improves antibody quality rather than reactogenicity or antigen expression. Together, these results define key parameters governing circRNA vaccine performance and provide a framework for optimising circRNA-based vaccines.

## Results

### RNA preparation quality is a key determinant of innate immune activation by circRNA

To investigate how circRNA purity influences innate immune responses in vivo, we used circRNA vaccines encoding two distinct antigens: the SARS-CoV-2 Wuhan spike protein receptor binding domain (RBD), expressed as a secreted trimer using a T4 foldon strategy^2^, and the nucleoprotein (NP) from influenza A virus (PR8; H1N1). All circRNA constructs were produced using intron–mediated autocatalytic circularisation and they incorporated the CVB3 IRES for cap-independent translation (Supplementary Figures 1 and 2).

During the initial phase of this project, we obtained two batches of circRNA for each antigen from the same supplier (Figure 1a). While the sequences in each batch were identical, the supplier’s quality control (by capillary electrophoresis) revealed significant differences in the levels of low molecular weight RNA species (Supplementary Table 1). In particular, the less refined preparations contained low molecular weight RNA species comprising 5.8% (RBD) and 10.6% (NP) of total RNA, likely arising from incomplete circularisation or splicing by-products, whereas these species were undetectable in the purer preparations. There is a well-established role for RNA contaminants of this type in driving innate immune activation by linear mRNA^21,36,37^. The differences in our batches therefore gave us a fortuitous opportunity to directly assess the contribution of low molecular weight RNA species to circRNA reactogenicity in vivo.

**Figure 1.**
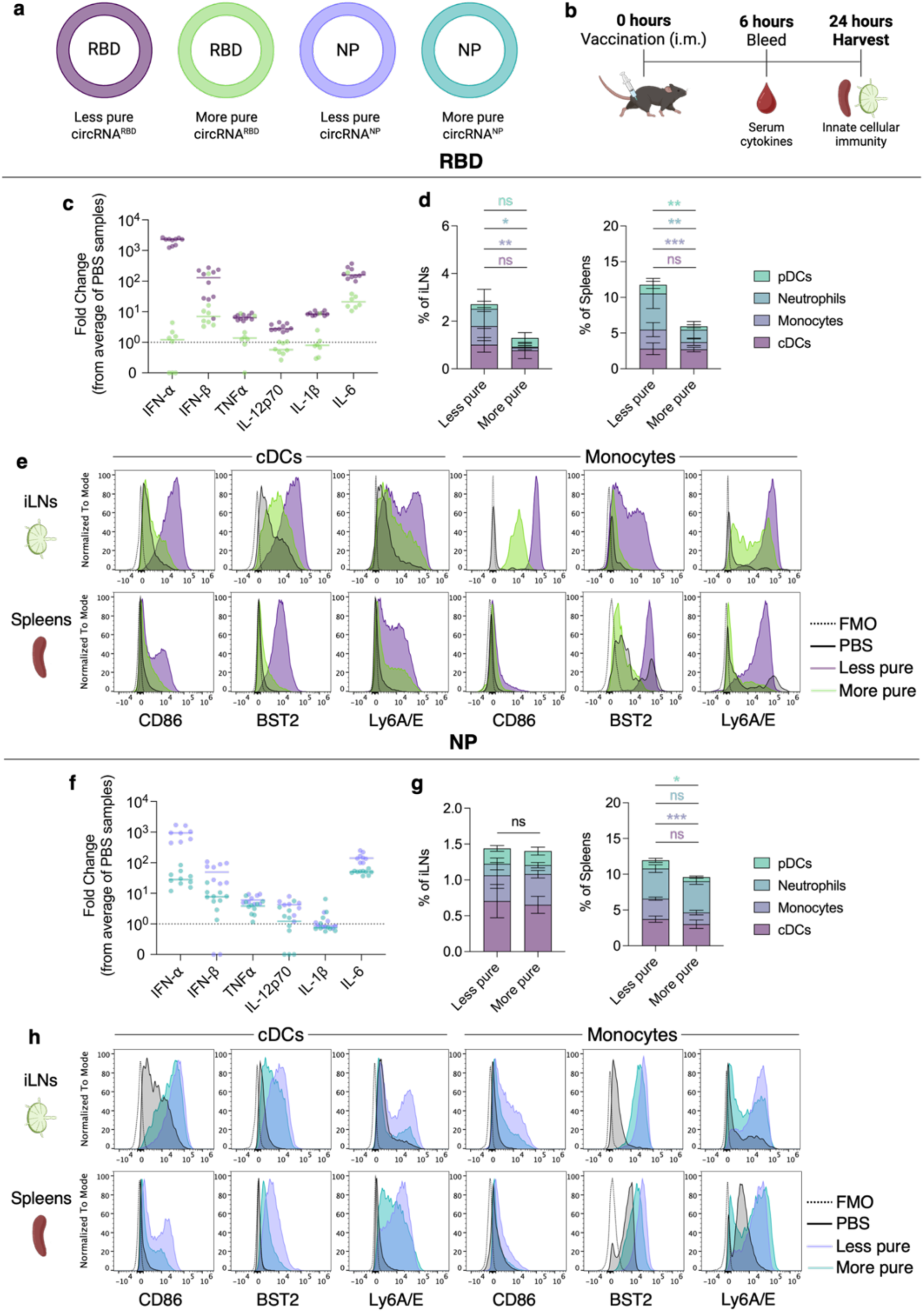
Innate immune activation by circRNA vaccines is determined by the quality of the RNA preparation. (**a**) Schematic of CircRNA constructs, emphasising the colour coding used throughout the figure. (**b**) C57BL/6J mice received 4 µg of LNP-encapsulated circRNA or PBS intramuscularly (i.m.); serum was collected by cheek bleed six hours post immunisation for cytokine analysis, and iLNs and spleens were harvested at 24 hours for assessment of innate immune cell populations. (**c** and **f**) Serum cytokine responses at six hours post-immunisation for (**c**) circRNA^RBD^ and (**f**) circRNA^NP^ vaccine groups. (**d** and **g**) Innate immune cell frequencies in iLNs and spleens for (**d**) circRNA^RBD^ and (**g**) circRNA^NP^ vaccine groups. (**e** and **h**) Representative histograms showing CD86, BST2, and Ly6A/E median fluorescence intensity (MFI) on cDCs and monocytes, in both iLNs and spleens for (**e**) circRNA^RBD^ and (**h**) circRNA^NP^ vaccine groups. Data represent *n* = 2 independent experiments (**c-f**) or *n* = 1 experiment (**g, h**) with *n* = 4-5 mice per group. Cytokine data (**c** and **f**) are expressed as median fold change relative to the PBS control group mean; dotted line indicates the PBS fold change reference. Bar graphs in (**d** and **g**) show mean ± SD; statistical comparisons were performed using an unpaired t-test, with ns = not significant; * P < 0.05; ** P < 0.01; *** P < 0.001. FMO, fluorescence minus one, RBD, receptor binding domain; NP, nucleoprotein.

**Figure 2.**
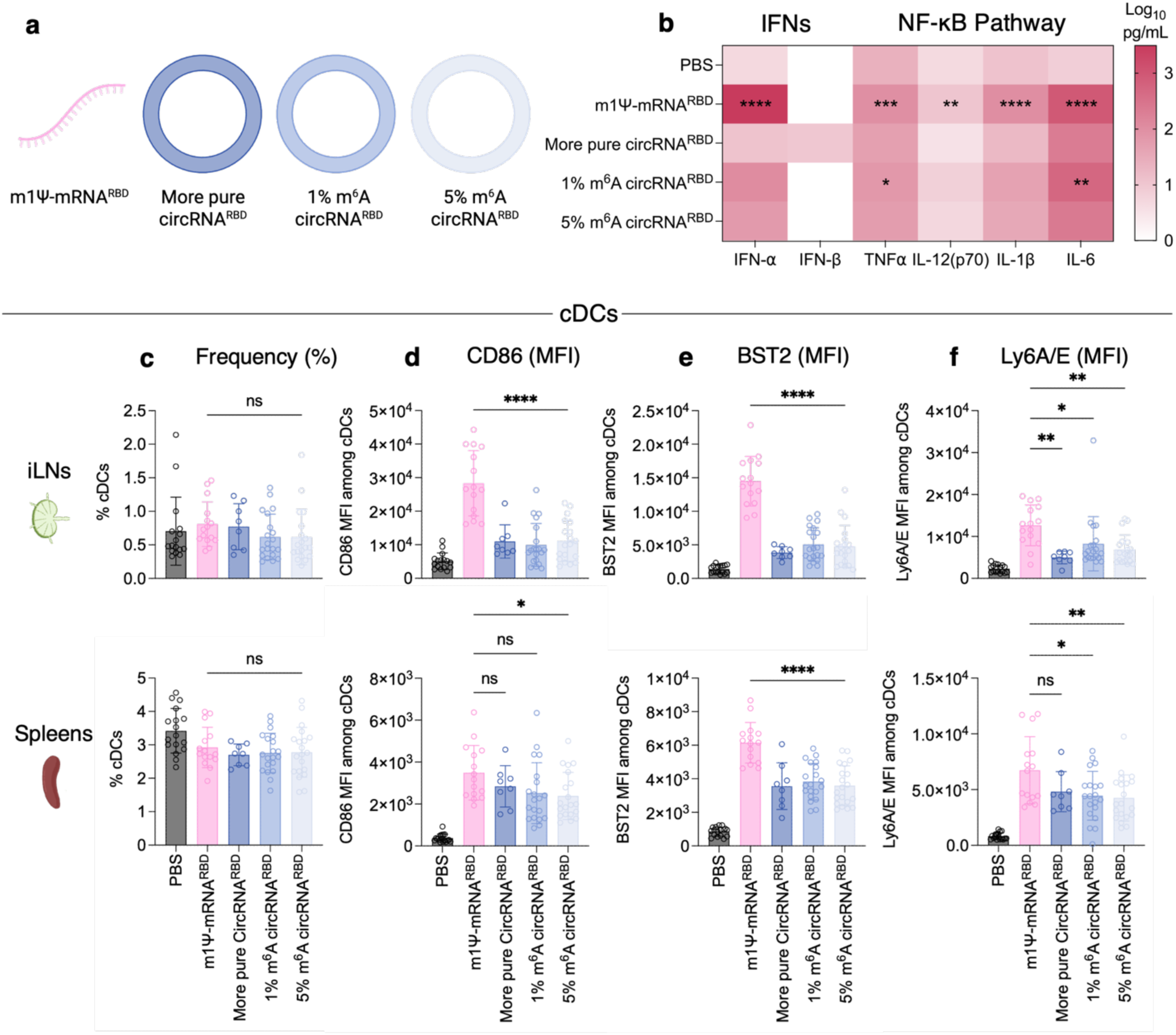
m^6^A modification does not further reduce reactogenicity of highly purified circRNA. (**a**) Schematic of RBD RNA vaccine constructs: mRNA incorporating 100% *N*^1^-methylpseudouridine (m1Ψ); unmodified circRNA (high preparation quality); and circRNA incorporating 1% or 5% m^6^A (**b**) Serum cytokine responses at 6 hours post-immunisation (log-transformed); statistical comparisons are relative to PBS negative controls. (**c-f**) Activation marker expression on cDCs in iLNs (top) and spleens (bottom); (**c**) cDC frequency (**d**) CD86 MFI (**e**) BST2 MFI (**f**) Ly6A/E MFI. Data represent *n* = 2-4 independent experiments with *n* = 3-5 mice per group, expressed as mean ± SD. Statistical comparisons were performed using a one-way ANOVA with Tukey’s multiple comparisons test, with ns = not significant; * P < 0.05; ** P < 0.01; *** P < 0.001; **** P < 0.0001.

C57BL/6 mice were immunised with circRNA delivered via lipid nanoparticle (LNP)-incorporating the ionisable lipid ALC-0315, and early innate immune responses were assessed. Serum cytokines were measured 6 hours post-immunisation to capture acute innate activation (Figure 1b). For the RBD antigen, less pure circRNA^RBD^ induced significantly higher levels of type I interferons (IFNs), particularly IFN-α, which was increased 1600-fold compared to the more pure circRNA^RBD^, which remained comparable to PBS controls (Figure 1c). In addition, proinflammatory cytokines associated with NF-κB activation, including tumour necrosis factor (TNF) α, interleukin (IL)-12p70, and IL-1β, were elevated in mice that received the less pure circRNA (2.3-, 4.7-, and 4.6-fold, respectively), with similar trends observed across additional cytokines and chemokines (Figure 1c and Supplementary Figure 3a). Notably, IL-6 was increased in the more pure circRNA^RBD^ group relative to PBS, consistent with known LNP-driven effects on IL-6 induction^38^.

To further characterise innate immune activation, we analysed immune cell populations in the draining inguinal lymph nodes (iLNs) and spleens 24 hours post-immunisation (Supplementary Figure 3b). Less pure circRNA^RBD^ increased the frequency of innate immune cells, including plasmacytoid dendritic cells (pDCs), neutrophils, and monocytes, relative to more pure circRNA^RBD^ (Figure 1d). While the overall frequency of conventional dendritic cells (cDCs) was similar between groups, cells from the less pure circRNA^RBD^ condition exhibited enhanced activation, as indicated by increased expression of the co-stimulatory molecule CD86 on both cDCs and monocytes (Figure 1e). In parallel, expression of interferon-stimulated surface markers BST2 and Ly6A/E were also elevated (Figure 1e), consistent with increased type I IFN signalling in response to less pure circRNA^RBD^ (Figure 1c).

Importantly, these findings were consistent across antigen systems. Less pure circRNA^NP^ similarly induced greater systemic cytokine responses and increased activation of innate immune cell populations compared to its purer counterpart (Figure 1f-h and Supplementary Figure 3c).

Together, these data demonstrate that circRNA purity is a major determinant of innate immune activation in vivo. The presence of low molecular weight RNA contaminants, potentially arising from circularisation by-products, drives increased type I interferon responses and innate cell activation, independent of the encoded antigen.

### m^6^A modification does not further reduce reactogenicity of purified circRNA

Having found that RNA purity is a primary determinant of circRNA-induced innate immune activation, next we sought to define the role of *N*^6^-methyladenosine (m^6^A) modification under conditions where circRNA is minimally immunostimulatory. Specifically, we asked whether m^6^A could further dampen residual reactogenicity, or whether its effects on circRNA function are independent of innate immune sensing.

To address this, we compared the more pure circRNA^RBD^ with constructs that were identical in sequence but incorporated 1% or 5% m^6^A (as a fraction of the total adenosine triphosphate pool) during in vitro transcription. As a benchmark, we included a linear mRNA encoding the same antigen containing m1Ψ (m1Ψ-mRNA^RBD^), representative of clinically approved mRNA vaccine formulations (Figure 2a, Supplementary Figure 4, Supplementary Table 1). All RNAs were formulated in ALC-0315-containing LNPs.

Assessment of systemic cytokine responses at 6 hours post-immunisation revealed that m1Ψ-mRNA^RBD^ induced the highest levels of proinflammatory cytokines and chemokines across most analytes tested (Figure 2b and Supplementary Figure 5a). In contrast, all circRNA^RBD^ formulations exhibited markedly reduced cytokine induction. Importantly, incorporation of m^6^A at either 1% or 5% did not further reduce cytokine responses relative to unmodified circRNA^RBD^.

We next examined innate immune cell recruitment and activation 24 hours post-immunisation. Vaccination with m1Ψ-mRNA^RBD^ resulted in increased frequencies of neutrophils, monocytes, and pDCs compared to all circRNA^RBD^ groups (Supplementary Figure 5b), whereas cDC frequencies were comparable between conditions (Figure 2c). Despite similar cellular composition, innate cell activation differed between circRNA and linear mRNA formats. cDCs and monocytes from circRNA^RBD^-immunised mice displayed significantly lower expression of the co-stimulatory molecules CD86 in iLNs, and interferon-stimulated markers BST2 and Ly6A/E in both iLNs and spleens compared with m1Ψ-mRNA^RBD^ (Figure 2d-f and Supplementary Figures 5c-e).

Consistent with the cytokine data, no differences in innate immune activation were observed between unmodified circRNA and m^6^A-modifed circRNA at either percentage. Together, these findings demonstrate that, in the context of highly purified circRNA, m^6^A modification does not further attenuate reactogenicity. Rather, circRNA exhibits a reduced immune profile compared to linear mRNA independent of m^6^A incorporation, suggesting that the lower reactogenicity of circRNA is driven by factors other than nucleotide modification.

### Purified circRNA supports greater antigen expression than linear mRNA, independent of m^6^A modification

Having established that RNA purity is a dominant determinant of circRNA reactogenicity and that m^6^A incorporation does not further modulate innate immune activation under these conditions, we next examined whether m^6^A influences circRNA antigen expression in vivo. Given that innate sensing can suppress translation, assessing antigen expression using highly purified, minimally immunostimulatory RNA is necessary to isolate the direct effects of m^6^A on translational efficiency.

We quantified antigen expression in vivo, after immunisation with purified circRNA^RBD^ constructs containing 0%, 1%, or 5% m^6^A, alongside m1Ψ-mRNA^RBD^ as a comparator. Muscle tissue from the injection site was harvested and homogenised at 24 and 48 hours post-immunisation, and RBD protein levels were measured by ELISA (Figure 3). At 24 hours, RBD expression by m1Ψ-mRNA^RBD^ averaged 94 ng/mg of tissue, which was 2.5- to 3.5-fold lower than RBD expression by all circRNA^RBD^ constructs (Figure 3). Notably, there were no significant differences in antigen expression between unmodified and m^6^A-modified circRNA at this timepoint, indicating that m^6^A does not influence protein output when RNA purity and innate immune activation are equivalent between groups.

**Figure 3.**
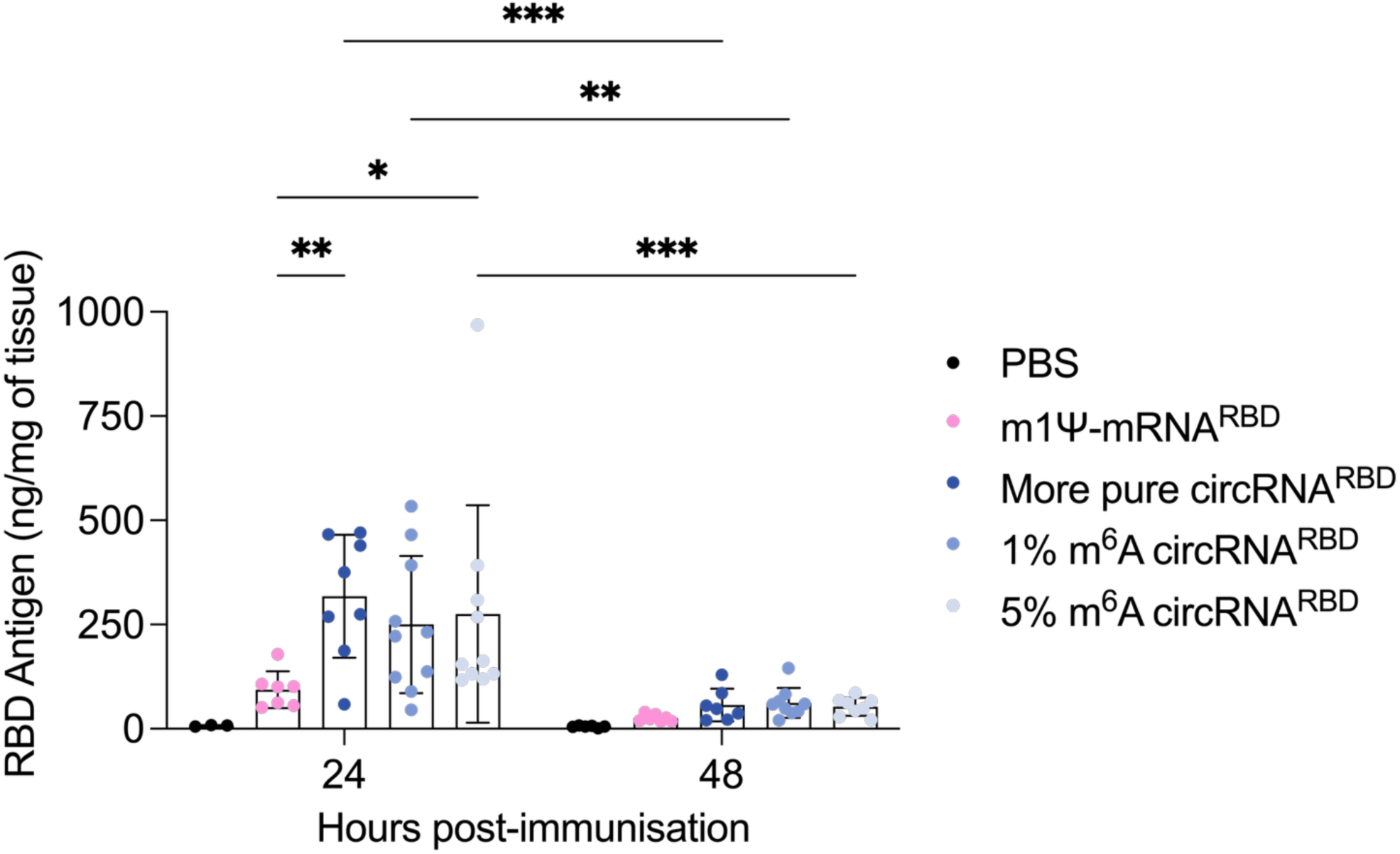
Purified circRNA drives greater antigen expression than linear mRNA, independent of m^6^A modification. C57BL/6J mice received 4 µg of LNP-encapsulated RNA vaccines i.m. Hindleg muscle was harvested and weighed at 24 and 48 hours post-immunisation. Clarified homogenates were assayed by ELISA using a capture anti-spike antibody and distinct detection anti-spike antibody, with amount of RBD antigen normalised to tissue weight. Data are a combination of *n* = 2 experiments with *n* = 3-5 mice per group, per experiment. Graph is depicted as mean ± SD. Two-way ANOVA with Tukey’s multiple comparisons test was performed, with * P < 0.05; ** P < 0.01; *** P < 0.001.

By 48 hours post-immunisation, antigen levels from all constructs (linear and circular) had declined to near-background levels, with no differences observed between m^6^A-modified and unmodified circRNA (Figure 3).

These findings suggest that purified circRNA drives greater antigen expression than linear mRNA in vivo, independent of m^6^A incorporation. In this context, the enhanced antigen expression observed with circRNA is most consistent with reduced innate immune activation rather than a direct effect of m^6^A on translational efficiency.

### CircRNA and linear mRNA elicit comparable adaptive immune magnitude, independent of innate immune reactogenicity

Next, we evaluated how RNA purity and m^6^A modification influence vaccine-induced adaptive immunity. We initially assessed immunogenicity using the SARS-CoV-2 RBD antigen, comparing circRNA and linear mRNA vaccines across formulations that differed in purity and nucleotide modification status (Figure 4a and Supplementary Table 1).

**Figure 4:**
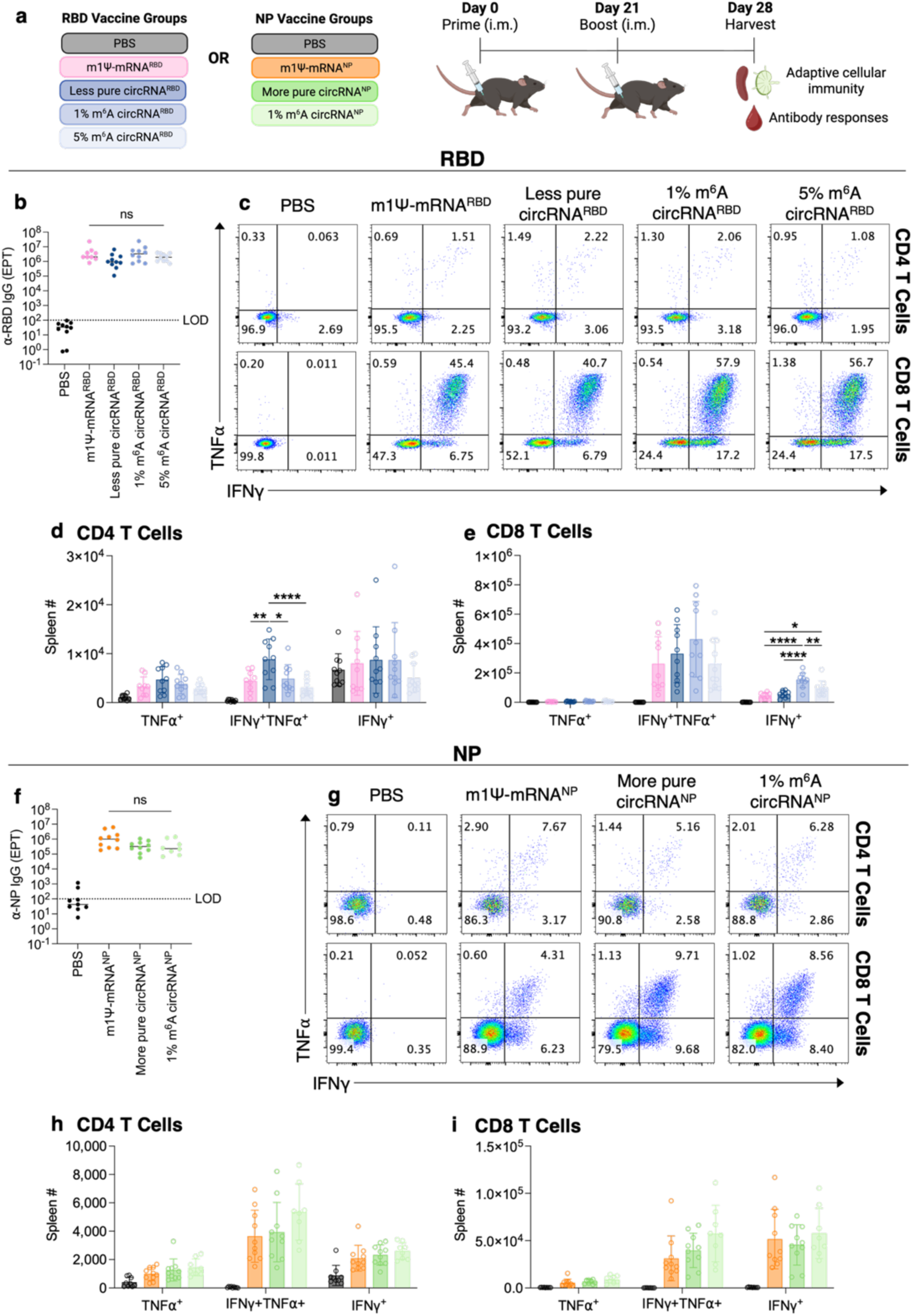
CircRNA and linear mRNA elicit comparable adaptive immune magnitude, independent of innate immune reactogenicity. (**a**) C57BL/6J mice received two 4 µg doses of LNP-encapsulated RNA vaccines i.m. three weeks apart. Spleens, iLNs, and serum were harvested one week after the second immunisation. (**b**) ⍺-Wuhan RBD IgG EPT in serum from RBD-vaccinated mice. (**c-e**) T cell responses in spleens from RBD-vaccinated mice: (**c**) representative flow plots showing TNF⍺ and IFNγ production by CD44^hi^ CD4 T cells (top panel) and CD44^hi^ CD8 T cells (bottom panel) following S1 peptide stimulation, and total numbers of cytokine-producing (**d**) CD44^hi^ CD4 T cells and (**e**) CD44^hi^ CD8 T cells. (**f**) ⍺-NP IgG EPTs in serum from NP-vaccinated mice. (**g-i**) T cell responses in spleen from NP-vaccinated mice: (**g**) representative flow plots showing TNF⍺ and IFNγ production by CD44^hi^ CD4 T cells (top panel) and CD44^hi^ CD8 T cells (bottom panel) following NP peptide stimulation, and total numbers of cytokine-producing (**h**) CD44^hi^ CD4 T cells and (**i**) CD44^hi^ CD8 T cells. Data represent *n* = 2 independent experiments with *n* = 4-5 mice per group, expressed as median (**b** and **f**) or mean ± SD (**d**, **e**, **h**, and **i**). Statistical comparisons were performed using a one-way ANOVA with Tukey’s multiple comparisons test, with ns = not significant; * P < 0.05; ** P < 0.01; **** P < 0.0001. IgG, immunoglobulin G; EPT, end point titre; LOD, limit of detection.

Despite the increased innate immune activation observed with less pure circRNA^RBD^ preparations (Figure 1c-e), we detected no significant differences in antigen-specific antibody titres between vaccine groups, as measured by ELISA quantifying antigen-specific IgG binding to recombinant Wuhan SARS-CoV-2 RBD protein (Figure 4b).

Antigen-specific T cell responses were assessed by intracellular cytokine staining following ex vivo stimulation of splenocytes with an S1 peptide pool, with flow cytometric quantification of IFNγ- and TNF⍺-producing CD4⁺ and CD8⁺ T cells (Supplementary Figure 6a). Overall, the magnitude of T cell responses was comparable across circRNA^RBD^ and m1Ψ-mRNA^RBD^ vaccines, irrespective of RNA purity or m^6^A modification (Figure 4c-e and Supplementary Figure 6b, c). However, we observed a modest reduction in a minor population of IFNγ single-producing T cells in the more reactogenic vaccine groups compared to less reactogenic formulations (Figure 4c, e and Supplementary Figure 6c).

Given that this population represented a small fraction of the total cytokine-producing T cells, which were dominated by IFNγ⁺TNF⍺⁺ double-producing cells, this difference is unlikely to have a major impact on overall T cell immunity. Nevertheless, it may suggest that the development of specific T cell functional subsets is sensitive to the level of innate immune activation.

To determine whether these observations were influenced by differences in reactogenicity, we next assessed responses in the influenza NP antigen system using highly purified circRNA constructs (Supplementary Figure 2, 7 and Supplementary Table 1), where innate immune activation was reduced. Consistent with the RBD dataset, NP-specific antibody titres were similar between unmodified circRNA^NP^, m^6^A-modified circRNA^NP^, and m1Ψ-mRNA^NP^ vaccines (Figure 4f). Antigen-specific T cell responses were also comparable in magnitude and functional profile, with no differences observed in cytokine-producing subsets, including IFNγ single-producing cells (Figure 4g-i and Supplementary Figure 6d, e). The absence of this difference under matched reactogenicity conditions indicates that the modest variation observed in the RBD system is likely attributable to differences in innate immune activation rather than RNA modality.

Across both antigen systems, incorporation of m^6^A at either 1% or 5% did not alter antibody titres or T cell responses. Together, these data demonstrate that RNA modality (linear or circular), reactogenicity, and m^6^A modification have minimal impact on the overall magnitude of adaptive immune responses in these models.

### m^6^A selectively enhances antibody quality, independent of the immune magnitude

While there were no significant differences in the antibody titres to any of our vaccines (Figure 4), it remained possible they were eliciting antibody responses of different qualities. To assess this, we employed functional assays that evaluated both antibody binding strength and neutralisation capacity. Antibody affinity was first assessed using a urea displacement ELISA, in which samples were treated with a high concentration of urea (6 M) to disrupt low-affinity interactions. This allowed us to quantify the fraction of the antibody pool that made strong enough interactions to survive this treatment. Across both NP and RBD antigen systems, incorporation of m^6^A into circRNA vaccines promoted the generation of ∼1.5-fold more high-affinity antibodies than unmodified circRNA and linear mRNA vaccines (Figure 5a, b). Notably, unmodified circRNA and linear mRNA elicited comparable affinity profiles independent of reactogenicity profiles, indicating that the enhancement in antibody binding strength was specific to m^6^A incorporation.

**Figure 5.**
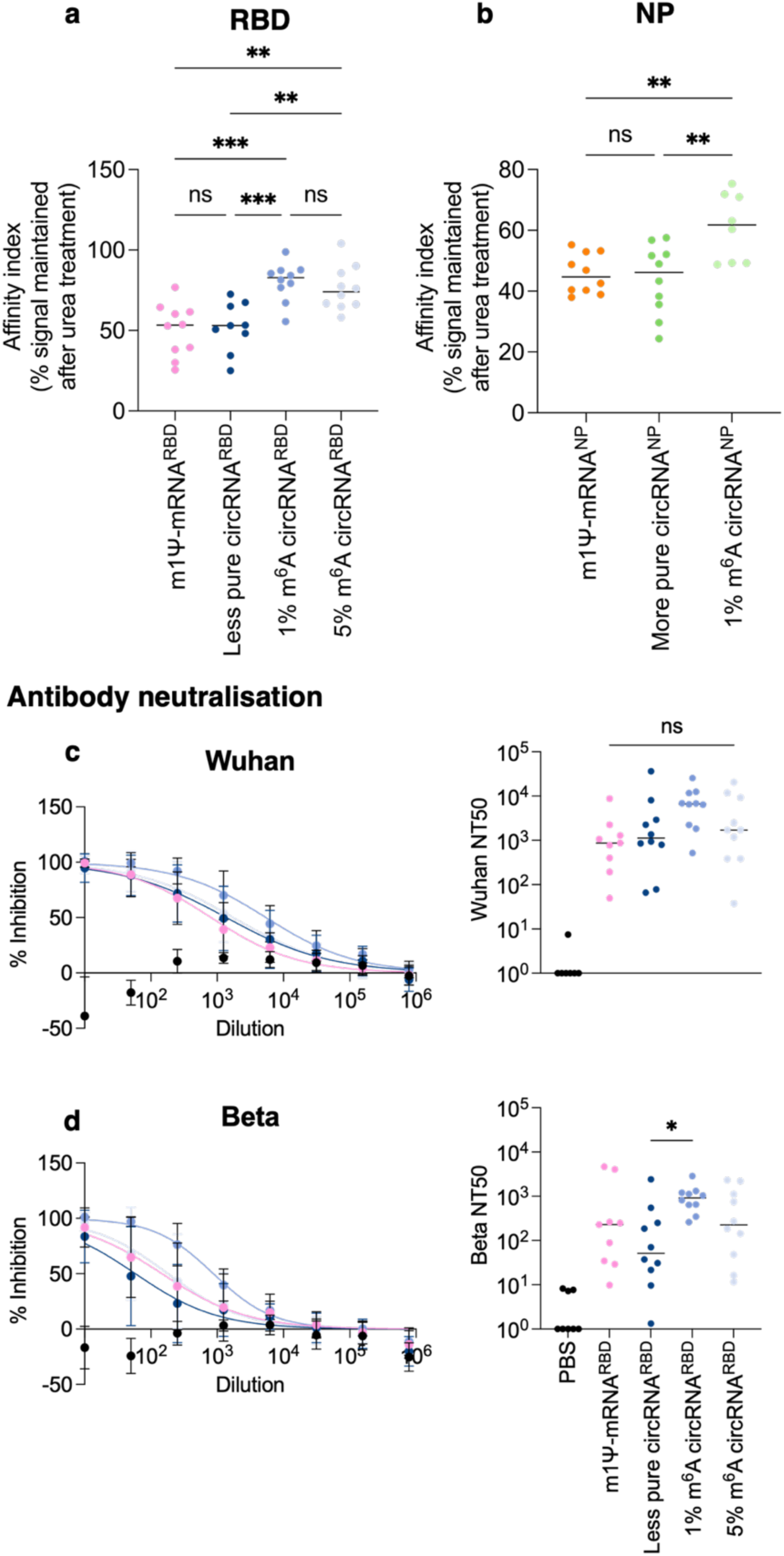
Incorporating m^6^A selectively enhances antibody quality. (**a** and **b**) Antibody affinity assessed by urea displacement ELISA on serum from mice immunised with (**a**) RBD or (**b**) NP vaccines. (**c** and **d**) Neutralisation of (**c**) Wuhan and (**d**) Beta pseudotyped viruses, shown as percent inhibition across serial dilutions (curve) and NT50 titres, measured in serum from RBD vaccinated mice. Data represent *n* = 2 experiments with *n* = 4-5 mice per group, expressed as median (**a** and **b**) or mean ± SD (**c** and **d**). Statistical comparisons were performed using a one-way ANOVA with Tukey’s multiple comparisons test, with ns = not significant; * P < 0.05; ** P < 0.01; *** P < 0.001.

To further evaluate antibody quality, we performed a pseudovirus neutralisation assay using viral particles pseudotyped with SARS-CoV-2 spike proteins. Neutralising activity was assessed against both the matched Wuhan strain (Figure 5c) and the related Beta variant (Figure 5d) to evaluate cross-reactive responses. Consistent with the affinity data, sera from mice immunised with m^6^A-modified circRNA^RBD^ exhibited the highest neutralising titres against the matched Wuhan strain (4.6-fold increase from m1Ψ-mRNA^RBD^ to 1% m^6^A circRNA^RBD^), although this did not reach statistical significance (P = 0.0618) (Figure 5c). In the cross-reactive Beta model, a similar trend toward increased neutralisation capacity was observed in the m^6^A-modified group relative to unmodified circRNA^RBD^ and m1Ψ-mRNA^RBD^ vaccines (Figure 5d).

Together, these findings demonstrate that incorporating m^6^A enhances the functional quality of the antibody response, as reflected by increased fractions of high-affinity binders and improved neutralisation capacity. Importantly, these effects were observed in the absence of changes in antibody magnitude or reactogenicity, indicating that m^6^A selectively improves antibody quality independent of its previously proposed role in modulating innate immune activation.

### CircRNA vaccination alters the anatomical distribution of germinal centre B cell responses

While m^6^A selectively enhanced antibody quality without affecting overall immune magnitude, these findings raised the possibility that qualitative differences in the humoral response may arise from alterations in B cell differentiation or organisation. We therefore examined antigen-specific B cell responses across lymphoid tissues to determine whether different RNA modalities influence the generation and anatomical distribution of germinal centre, memory and plasma B cell populations. Antigen-specific B cells were identified using fluorescent tetramers generated from biotinylated RBD or NP proteins complexed with fluorophore-conjugated streptavidin. To ensure specificity, a dual-labelling strategy was employed in which each antigen was labelled with two distinct fluorophores, and only B cells binding both tetramers were included in downstream analyses. Non-specific binders were excluded using a decoy tetramer generated from biotinylated 4-hydroxy-3-nitrophenylacetyl-bovine serum albumin (NP-BSA); B cells binding the decoy tetramer were excluded from all downstream analysis (Supplementary Figure 8a).

B cell subsets were first defined as B220⁺IgD⁻ cells, with germinal centre (GC) B cells identified as GL7⁺CD38⁻CD95⁺Bcl6⁺and memory B cells as GL7⁻CD38⁺. Within each of these subsets, the frequency of antigen-specific cells was then determined by tetramer binding. Plasma cells were identified as B220⁻CD138⁺IRF4⁺cells (Supplementary Figure 8a). Antigen-specific T follicular helper (T_FH_) cells were identified using NP-MHC II tetramers to directly enumerate Bcl6⁺PD1⁺CD4⁺ T cells recognising a dominant NP epitope, allowing assessment of antigen-specific T cell help within germinal centres (Supplementary Figure 8b). These populations were assessed in both iLNs and spleens following vaccination.

Responses in the iLNs were broadly comparable across vaccine groups, with no significant differences in GC B cells, memory B cells, plasma cells, or T_FH_ cells, between RNA modalities, for either antigen systems (Figure 6a-f and Supplementary Figure 8c-h). In contrast, analysis of the spleen revealed a consistent and specific reduction in GC B cell responses following circRNA vaccination (Figure 6g-l). While total GC B cell frequencies were unchanged (Supplementary Figure 8i, l), antigen-specific GC B cells were markedly reduced across both RBD and NP antigen systems (Figure 6g, j). Splenic memory B cells and plasma cells were largely preserved (Figure 6i, l and Supplementary Figure 8j, m), with RBD-specific memory B cells unchanged between groups (Figure 6h) and only a modest reduction in NP-specific memory B cells observed with circRNA vaccination (Figure 6k). To further interrogate the splenic GC defect, we examined antigen-specific T_FH_ cells in the NP system, where tetramer reagents permitted enumeration of the antigen-specific response. Whereas NP^+^ T_FH_ cells were intact in the iLNs (Figure 6m), they were markedly reduced in the spleen following circRNA^NP^ vaccination (Figure 6n), with total T_FH_ cell frequencies being unaffected (Supplementary Figure 8k, n). Together, the parallel reduction in both antigen-specific GC B cells and antigen-specific T_FH_ cells in the spleen, with responses intact in the iLN, suggest a compartment-specific failure to establish a splenic GC response following circRNA vaccination, which was independent of m^6^A modification.

**Figure 6.**
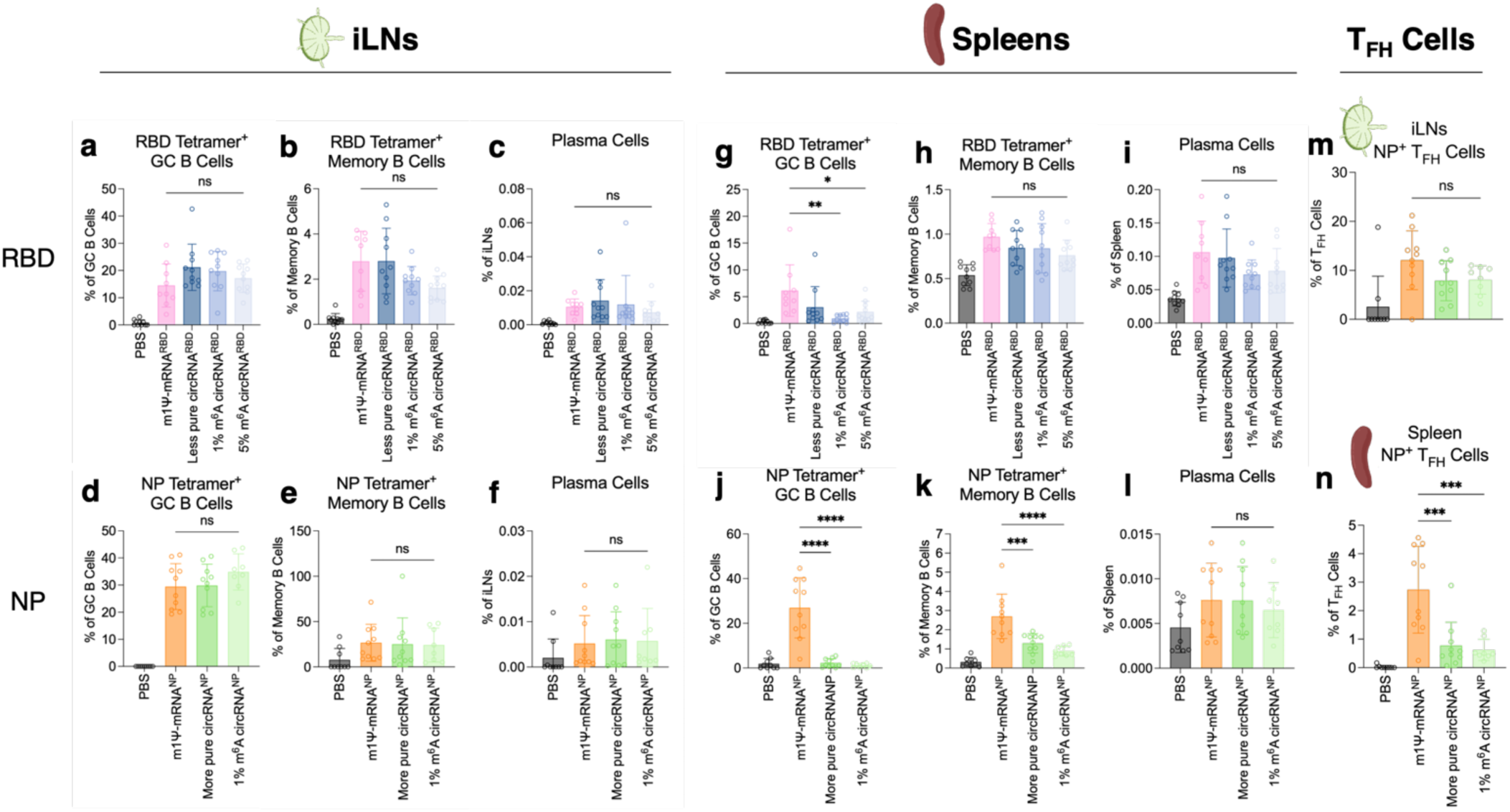
CircRNA vaccination fails to establish splenic germinal centre responses while preserving lymph node immunity. (**a–c**) B cell responses in iLNs of RBD-vaccinated mice: frequency of RBD-specific cells within (**a**) GC B cells and (**b**) memory B cells, and (**c**) total plasma cell frequency. (**d–f**) Corresponding populations in the iLNs of NP-vaccinated mice: frequency of NP-specific cells within (**d**) GC B cells and (**e**) memory B cells, and (**f**) total plasma cell frequency. (**g–i**) B cell responses in spleens of RBD-vaccinated mice: frequency of RBD-specific cells within (**g**) GC B cells and (**h**) memory B cells, and (**i**) total plasma cell frequency. (**j–l**) Corresponding populations in the spleens of NP-vaccinated mice: frequency of NP-specific cells within (**j**) GC B cells and (**k**) memory B cells, and (**l**) total plasma cell frequency. (**m** and **n**) Frequency of NP-specific T_FH_ cells within the total T_FH_ pool, identified using NP-MHC II tetramers enumerating Bcl6⁺PD1⁺CD4⁺ T cells, in (**m**) iLNs and (**n**) spleens. GC B cells and memory B cells were defined as B220⁺IgD⁻GL7⁺CD38⁻ and B220⁺IgD⁻GL7⁻CD38⁺ respectively; antigen-specificity was determined by dual-fluorophore tetramer binding within each subset. Plasma cells were identified independently as B220⁻CD138⁺IRF4⁺. Non-specific tetramer binders were excluded using a 4-hydroxy-3-nitrophenylacetyl bovine serum albumin (NP-BSA) decoy tetramer. Data represent *n* = 2 independent experiments with *n* = 4–5 mice per group, expressed as mean ± SD. Statistical comparisons were performed using a one-way ANOVA with Tukey’s multiple comparisons test, with ns = not significant, * P < 0.05; ** P < 0.01; *** P < 0.001 **** P < 0.0001.

## Discussion

CircRNA is emerging as a promising next-generation vaccine strategy. However, the field has lacked systematic in vivo comparisons that disentangle the relative contributions of RNA purity, nucleotide modification, antigen expression and innate immune activation to vaccine-induced immunity. This has led to an ongoing debate as to whether circRNA is intrinsically less reactogenic than linear mRNA, and whether this reduced reactogenicity directly translates into improved vaccine efficacy^9^. In this study, we addressed these questions through a comprehensive in vivo analysis across two antigen systems, integrating measurements of innate immune activation, antigen expression, adaptive immune magnitude, antibody quality and anatomical localisation of B cell responses.

A central finding from this work is that the quality of circRNA preparations, specifically in the presence of residual low-molecular-weight RNA species, is a major determinant of innate immune activation in vivo. Less pure circRNA preparations induced elevated type I interferon responses and increased activation of innate immune cell populations, whereas highly purified circRNA exhibited minimal reactogenicity. These findings are consistent with prior work demonstrating that RNA byproducts generated during in vitro transcription and circularisation, such as linear precursors, excised introns and nicked RNA species, can form immunostimulatory structures including double-stranded RNA and 5′ triphosphate-containing RNA capable of activating RIG-I and MDA5^1,30,33,39^. Similar principles have been established in the mRNA field, where dsRNA contaminants suppress translation and increase inflammatory signalling, leading to the adoption of purification strategies such as HPLC^21,36,37^.

Our findings also help to reconcile conflicting reports regarding circRNA reactogenicity. Early studies suggested that circRNA generated using group I intron-based methods could activate innate sensing pathways due to the scar sequence left after splicing^8,40^. On the other hand, subsequent work demonstrated that purified circRNA exhibits reduced reactogenicity and prolonged translation. More recent studies have further suggested that RNA preparation quality, rather than scar sequence alone, is the dominant determinant of circRNA immunogenicity^1,30^. Our in vivo data support this interpretation, showing that innate activation tracked directly with preparation quality across two independent antigen systems. Importantly, when circRNA was highly refined, m^6^A incorporation did not further reduce reactogenicity, indicating that low innate activation is primarily achieved through RNA preparation quality rather than nucleotide modification.

Despite these differences in innate immune activation (Figure 2), reactogenicity had limited impact on the overall magnitude of adaptive immunity (e.g. Figure 4b and 4f). This finding contrasts with the classic model for linear mRNA, in which reducing innate sensing enhances protein expression and improves immunogenicity^19,41^, though the relationship between nucleotide modification and adaptive immune magnitude in linear mRNA systems has itself proven more nuanced in practice^42–44^. The basis for this decoupling for circRNA is not fully resolved by the current data and likely reflects the multifactorial nature of vaccine-induced immunity in vivo. It is worth noting that lipid nanoparticle delivery systems themselves are known to provide potent adjuvant signals that promote T_FH_ and humoral responses^38^, which may contribute to the robustness of adaptive immunity regardless of differences in the intrinsic reactogenicity of the RNA species being tested. Fully dissecting the relative contributions of these parameters will require further investigation.

An important and unexpected finding was that m^6^A modification enhanced antibody quality without affecting reactogenicity, antigen expression or overall immune magnitude. Incorporating m^6^A into our circRNA vaccines increased the fraction of high-affinity antibodies that were produced, and also improved neutralisation capacity, particularly against matched viral strains. This indicates that m^6^A influences the quality of humoral immunity through mechanisms distinct from its proposed role in reducing innate immune sensing. Mechanistically, m^6^A has been shown to regulate RNA translation and stability through recruitment of reader proteins such as YTHDF1, which enhances translation via interaction with eIF3^45,46^. In addition, m^6^A can promote cap-independent translation through IRES-like mechanisms, particularly when located within specific sequence contexts such as the 5′ UTR or internal regions^47^. However, in our system, random incorporation of m^6^A during in vitro transcription will have resulted in heterogeneous positioning, which may explain why no increase in overall antigen expression was observed.

Another key finding from this study is that circRNA vaccination selectively lacks splenic germinal centre responses while preserving comparable responses in the draining LN. This was characterised by consistent reductions in antigen-specific GC B cells and T_FH_ cells in the spleen across both antigen systems. Notably, this compartment-specific pattern was independent of m^6^A modification, suggesting this finding is a feature of the circRNA modality rather than nucleotide modification status.

The mechanistic basis for this tissue-specific effect remains to be determined. However, the selectivity for GC B cells rather than memory or plasma cell compartments is notable. Unlike memory B cells and plasma cells, which persist independently of ongoing antigen engagement and redistribute between lymphoid compartments via the circulation, GC reactions are uniquely dependent on sustained local antigen availability for their maintenance^48^. One parameter that could differ between RNA modalities and between lymphoid compartments is antigen availability over time. Sustained antigen delivery has been shown to be a key determinant of GC magnitude^49^, and circRNA and linear mRNA may differ in the duration or tissue distribution of antigen expression due to differences in RNA stability or translation kinetics^50^. We do not directly demonstrate this here, and the precise mechanism remains an open question, but the compartment-specific pattern we observe is consistent with a model in which local antigen availability differs between tissues following circRNA vaccination.

Together, these findings have important implications for circRNA vaccine design. They demonstrate that reduced reactogenicity alone is not sufficient to enhance adaptive immune magnitude, and that circRNA vaccine performance is shaped by multiple independent parameters. RNA purity is a dominant determinant of innate immune activation; m^6^A modification enhances antibody quality without affecting magnitude or reactogenicity; and the choice of RNA modality influences the anatomical distribution of antigen-specific B cell responses. This highlights the need to evaluate RNA vaccines not only in terms of expression and inflammation, but also in terms of immune quality and tissue-specific responses.

This study has illuminated several avenues for future research. While we demonstrated a clear relationship between circRNA purity and innate immune activation, sequencing and biochemical analysis will be required to precisely define the contaminating species responsible for the effect. In addition, broader mapping of antigen production across different cell types will likely be critical for understanding the mechanism underlying tissue-specific germinal centre responses. Finally, the mechanism by which m^6^A enhances antibody quality remain to be elucidated and will require targeted studies of antigen presentation, B cell selection and T_FH_ cell responses.

In summary, this study provides a systematic in vivo framework for understanding how circRNA vaccine design parameters influence immune outcomes. We show that highly purified circRNA is minimally reactogenic, but that reduced innate activation does not necessarily enhance adaptive immune magnitude. We further demonstrate that m^6^A modification selectively improves antibody quality, and that circRNA vaccination alters the anatomical distribution of antigen-specific B cell responses. These findings define key parameters for the rational design of circRNA vaccines and highlight the importance of considering not only immune magnitude, but also immune quality and spatial organisation.

## Methods

### Expression and purification of AviTag RBD protein

Fc- and Avi-tagged RBD protein containing an HRV-3C protease cleavage site were expressed in Expi293F suspension cells (Gibco) using the ExpiFectamine method according to the manufacturer’s protocol. Cells were cultured at 37 °C with 9% CO_2_, 120 rpm for five days. CaptivA PriMAB rProtein A Affinity Resin (Repligen) was saturated with Fc-tagged proteins by running conditioned media through a column. Resin was washed with high salt buffer (50 mM Tris pH 8.0, 450 mM NaCl), then equilibrated with low salt buffer (50 mM Tris pH 8.0, 150 mM NaCl). Resin was incubated in low salt buffer containing 1 mM DTT and 10 μg/mL human rhinovirus (HRV)-3C protease in low salt buffer was added and left overnight to cleave proteins, which all contained an HRV-3C protease cleavage site. Protein was then eluted and concentrated to 1-5 mg/mL using 10 kDa centrifugal filters at 1,800*g*, 4 °C. A NanoDrop One instrument (Thermo Fisher Scientific) was used for determining concentrations, with extinction coefficients obtained using ProtParam tool^51^. AviTag RBD proteins were biotinylated using the Sigma Aldrich Enzymatic Protein Biotinylation kit, and biotinylation was confirmed through ELISA.

### Expression and purification of AviTag NP protein

NP containing both an AviTag and a His_6_ tag was expressed in *E. coli* AVB101 (Avidity) which contains an IPTG-inducible *birA* gene to allow in vivo protein biotinylation. Cultures were grown in TYH medium supplemented with 0.5% glucose (220 rpm, 37 °C) until an OD_600nm_ of 0.4-0.6 was reached. Biotin (50 µM) and IPTG (0.5 mM) were added, and the culture was incubated overnight (180 rpm, 20 °C). Cells were then pelleted by centrifugation (30,000*g*, 30 mins, 4 °C) and lysed by sonication in binding buffer (40 mM Tris, 300 mM NaCl, 20 mM imidazole, pH 7.6). NP was subsequently purified from the clarified cell lysate using a HisTrap HP column (Cytiva) with a gradient elution of binding buffer containing 500 mM imidazole. The affinity purified NP was further polished using a HiLoad 16/600 Superdex 200 pg column. The SEC-purified NP was concentrated using 10 kDa centrifugal filters (4,000*g*, 4 °C) and protein concentration determined using a Pierce BCA Protein Assay Kit (Thermo Fisher Scientific). Protein was used immediately or aliquoted, flash frozen and stored at -80 °C for later use. Biotinylation was confirmed through ELISA.

### RNA

All RNA constructs used in this study, including their sequences and design elements, were synthesised by GenScript (Piscataway, NJ, USA). Quality controls on RNA purity were carried out by the manufacturer using capillary electrophoresis and are summarised in Supplementary Table 1.

### Lipid nanoparticle encapsulation of RNA

RNA-LNPs were formulated using a NanoAssemblr Ignite (Precision Nanosystems), as described previously^52^. The lipid composition was based on the ALC-0315 formulation used in the Pfizer COVID-19 vaccine. RNA was encapsulated at a final concentration of 100 μg/mL, and LNP suspensions were stored at 4 °C until use.

### Mice

C57BL/6J mice were bred and housed under specific pathogen-free conditions at the Malaghan Institute of Medical Research Biomedical Research Unit (Wellington, New Zealand). C57BL/6J mice were originally obtained from The Jackson Laboratory. Sex-matched male and female mice between 8-12 weeks of age were used for experiments. All experiments were carried out in accordance with the guidelines stated by the Victoria University of Wellington Animal Ethics Committee (30841: Assessing Immunogenicity of Novel COVID-19 Vaccines) and followed the Code of Ethical Conduct for the Manipulation of Animals.

### Immunisations

LNP-RNA vaccines were prepared with sterile PBS such that each mouse would receive 4 μg at 100 μL. Ketamine/xylazine-sedated mice were intramuscularly injected with 50 µL of prepared vaccine on both legs, each injection containing equal amounts of LNP-RNA. The dose refers to the total amount of antigen injected per mouse.

### Tissue preparation

Following sacrifice, blood, spleens and both iLNs were collected for analysis. Cardiac punctures were performed to obtain blood and placed into a Microvette tube (Sarstedt). These were centrifuged at 10,000*g* for 5 minutes, then serum was aliquoted and stored at -80 °C. To prepare single cell suspensions of splenocytes, spleens were mashed through 70 μm cell strainers (Falcon) and washed through with Iscove’s Modified Dulbecco’s Medium (IMDM) (Gibco), then centrifuged for 10 minutes at 250*g*, 4 °C. Cell pellets were resuspended in Puregene Red Blood Cell Lysis Solution (Qiagen) before centrifugation for 4 minutes at 400*g*, 4 °C. The cell pellet was refiltered through a 70 μm cell strainer in IMDM, then counted and centrifuged for 10 minutes at 250*g*, 4 °C. Cell pellets were resuspended to 2×10^7^ cells/mL in R10 media (RPMI [Rosewell Park Memorial Institute] 1640 medium with 10% FCS [foetal calf serum, Gibco]) and plated for flow cytometry staining. When investigating DCs, spleens were digested with Liberase TL Research Grade (Roche) and DNase I (Roche) in IMDM for 25 minutes at 37 °C. Digested splenocytes were then processed as previously mentioned, but following red blood cell lysis and centrifugation, the cell pellet was resuspended in IMDM and plated for flow cytometry staining. To prepare single cell suspensions of iLNs, tissues were mashed through 70 μm cell strainers and washed through with IMDM, then centrifuged for 10 minutes at 250*g*, 4 °C. Cell pellets were then resuspended in IMDM and plated for flow cytometry staining. When investigating DCs, iLNs were teased apart using two 18G needles before being digested with Liberase TL Research Grade and DNase I in IMDM for 25 minutes at 37 °C. Digested iLNs were then processed and plated for flow cytometry staining as previously mentioned.

### Intracellular cytokine staining (ICS)

For intracellular cytokine staining (ICS), splenocytes were plated with PepMix SARS-CoV-2 (S-RBD) or Influenza NP (H1N1 PR8) (JPT Peptide Technologies) in 10 µg/mL Brefeldin A (Abcam) according to the manufacturer’s protocol. The samples were incubated at 37 °C for 6 hours, then kept at 4 °C overnight to be stained the following day.

### B cell tetramer formation

Biotinylated NP-BSA, Wuhan RBD, and NP proteins were prepared as separate reactions to add fluorescently labelled streptavidin (SAV). Fluorescently labelled SAV was added at a 1:20 protein to SAV ratio, where a fifth was added to the protein at a time, with 30-minute incubations between additions, on ice. Once all the SAV was incorporated, 4 mM free biotin (Sigma) in PBS was added at a 1:1 volume ratio for 1 hour, to saturate free SAV. Tetramers were centrifuged for 10 minutes at 18,000*g*, 4 °C, then combined to make a tetramer cocktail. The decoy biotinylated NP-BSA formed tetramers with SAV BV711 to first exclude any non-specific binding B cells. Biotinylated Wuhan RBD protein formed tetramers with SAV PE-CF594 and SAV PE-Cy5, and biotinylated NP protein formed tetramers with SAV PE and SAV APC for the dual-labelling strategy to ensure specificity.

### Flow cytometry staining

Single cell suspensions were stained for viability with Zombie NIR (BioLegend) or BD Horizon Fixable Viability Stain 700 (BD Biosciences), before incubation with anti-mouse CD16/32 (clone 2.4G2), to block any non-specific antibody binding. To stain for NP⁺CD4⁺ TFH cells, samples were incubated for 1 hour at RT with a cocktail containing an APC-labelled NP MHC-II tetramer (epitope: QVYSLIRPNENPAHK I-A(b)). To stain for surface markers, cells were incubated for 15 minutes at 4 °C with cocktails of fluorescent antibodies specific for: IgD (11-26c.2a), CD79b (HM79-12), CD44 (IM7), CD95 (Jo2), GL7 (GL-7), CD4 (GK1.5), B220 (RA3-6B2), TCRβ (H57-597), PD-1 (29F.1A12), CD19 (6D5), CD138 (281-2), CD38 (90), CD8⍺ (53-6.7), CD64 (X54-5/7.1, H1.2F3), CD86 (GL1), CD11b (M1/70), CD11c (N418), Ly6C (HK1.4), Ly6G (1A8), BST2 (927), MHC II (2G9), and Ly6A/E (D7). Prior to tetramer staining, cells were blocked with 20% FCS on ice for one hour. The tetramer cocktail was then applied and incubated on ice for 30 minutes. To stain for intranuclear transcription factors, cells were fixed and permeabilised using the eBioscience FoxP3/Transcription Factor Staining Buffer Set (Invitrogen) according to the manufacturer’s protocol. Cells were then incubated for 1 hour at RT with cocktails of fluorescent antibodies specific for: IRF4 (3E4), Bcl6 (K112-91), IFNγ (XMG1.2), or TNF⍺ (MP6-XT22). If no intranuclear staining was required, cells were fixed using formalin solution (Sigma-Aldrich) and incubated for 20 minutes at 4 °C. Data were acquired on a Cytek Aurora (Cytek Biosciences) and analysed using FlowJo version 10.10.0 (BD Biosciences).

### Conventional ELISA for detecting serum IgG_T_

To detect Wuhan RBD- or NP-specific IgG total (IgG_T_) in serum, Nunc MaxiSorp 96-Well Microplates (Thermo Fisher Scientific) were incubated overnight at 4 °C with 2 μg/mL Wuhan RBD or NP protein in bicarbonate buffer. Plates were blocked with 10% FCS to prevent non-specific binding before adding serially diluted serum and incubating for 2 hours. Samples were then incubated with goat ⍺-mouse IgG_T_-HRP detection antibody (Invitrogen) before developing reactions with tetramethylbenzidine (TMB; BD Biosciences), which were subsequently terminated using 2 M H_2_SO_4_ then immediately read at 450 nm using a plate reader (VICTOR Nivo, PerkinElmer). After each incubation, plates were washed 5× with 0.05% TWEEN 20 (Sigma-Aldrich) in PBS. Endpoint titres were calculated using the t-statistic method as previously described^53^.

### Urea displacement assay

To evaluate antibody affinity, the conventional ELISA protocol was followed as described above. Following the 2-hour sample incubation, an additional incubation with 6 M urea (Sigma-Aldrich) or PBS was conducted for 30 minutes at 37 °C. The remaining steps were carried out as described above. Affinity index was calculated as previously described^54^.

### RBD protein expression ELISA

To process tissue for RBD protein quantification, hind leg muscle tissue was harvested from mice at various timepoints post-vaccination. Harvested muscles were snap-frozen in liquid nitrogen, then weighed for normalisation. A volume of 300 µL/100 mg of protease mix (5 mM EDTA and 1× Halt Protease and Phosphatase Inhibitor Cocktail [Thermo Fisher Scientific] in PBS) was added to each tissue, then homogenised using a T 25 Basic Ultra-Turrax homogeniser (IKA) for 10-20 seconds. Tissue homogenates were centrifuged at 15,000*g* for 10 minutes at 4 °C, and the supernatant transferred to a new tube and centrifuged for another 10 minutes at 15,000*g*. After the second centrifugation, tissue homogenates were aliquoted and stored at -80 °C until ELISA.

To quantify RBD protein expression in the muscle tissue, Nunc MaxiSorp 96-Well Microplates (Thermo Fisher Scientific) were incubated overnight at 4 °C with 1 μg/mL rabbit anti-spike neutralising antibody (SinoBiological, Cat: 40592-R001) in bicarbonate buffer. Plates were blocked with 20% FCS to prevent non-specific binding before adding neat tissue homogenates. Samples (along with RBD protein of a known concentration to generate a standard curve) were then incubated for two hours with 0.122 µg/mL mouse anti-spike neutralising antibody (SinoBiological, Cat: 40591-MM43) diluted in 20% FCS. Samples were then incubated with goat ⍺-mouse IgG_T_-HRP detection antibody before developing reactions with TMB, which were subsequently terminated using 2 M H_2_SO_4_ then immediately read at 450 nm using a plate reader. After each incubation, plates were washed 5× with 0.05% TWEEN 20 (Sigma-Aldrich) in PBS. Raw OD450 nm readings were interpolated with the standard curve using the Sigmoidal, 4PL, X is concentration equation in Prism to obtain sample RBD concentrations (µg/mL). These were then converted to ng/mg of tissue by normalising to the tissue weights collected post-harvest.

### LEGENDplex

To quantify cytokines in serum obtained from mice, the LEGENDplex Mouse Anti-Virus Response Panel (BioLegend) and the associated LEGENDplex Data Analysis Software was used. Serum samples were diluted 1:8, and the assay undertaken according to the manufacturer’s protocol. Data were acquired on a Cytek Aurora (Cytek Biosciences).

### Generation of SARS-CoV-2 spike pseudotyped lentiviruses

Pseudotyped lentiviral particles, incorporating the spike glycoprotein from SARS-CoV-2 with 19 amino acids removed from the carboxy terminal, were generated as described previously^55^. Briefly, HEK293FT cells were seeded in 6-well plates (4 × 10^5^ cells/well, in 3 mL of DMEM [Dulbecco’s Modified Eagle Medium, Gibco], 10% FBS, 500 µg/mL of geneticin, 2 µg/mL of puromycin) and incubated at 37 °C, 5% CO_2_ for 24 hours. Cells were then co-transfected, using FuGENE HD Transfection Reagent (Promega), with the following plasmids: (i) 1 µg of pHAGE-CMV-Luc2-IRES-ZsGreen-W, (ii) 0.22 µg of HDM-Hgpm2, (iii) 0.22 µg of pRC-CMV-Rev1b, (iv) 0.22 µg of HDM-tat1b, and (v) 0.37 µg of a plasmid containing a codon optimised SARS-CoV-2 spike protein. In addition to the plasmid expressing the spike protein from the original ancestral isolate (pcDNA3.1_spike_del19, a gift from R. DeFrancesco, plasmid #155297, Addgene), a series of plasmids were synthesised to generate pseudotyped viral particles containing codon optimised versions of the spike glycoprotein from ancestral (Wuhan) or B.1.351 (Beta) SARS-CoV-2 variants, using a pcDNA3.1 backbone (Gene Universal). Cell culture media was changed 24 hours after transfection and cell-free supernatant collected 48 hours post-transfection, filtered through a 0.45 µm syringe filter (Interlab) and 3 mL aliquots stored at -80 °C until future use. The titre of the SARS-CoV-2 spike pseudotyped lentiviruses was determined by infecting HEK293 cells stably expressing human ACE2 (HEK293/ACE2 cells, obtained from Dr. John Taylor, University of Auckland).

### Pseudotyped virus neutralisation assay

The ability of mouse serum samples to neutralise SARS-CoV-2-spike-mediated entry was determined as described previously^55^. Briefly, HEK293/ACE2 cells were seeded in poly-D-lysine (Gibco)-coated, white-walled, 96-well plates (25,000 cells/well) and incubated at 37°C, 5% CO_2_ for 24 hours. Serum samples collected from immunised mice were heat-treated at 56 °C for 30 minutes, diluted with cell culture medium plus 7.5 µg/mL polybrene (Sigma-Aldrich; 1:10, then 1:5 serial dilutions), mixed with a suspension of the SARS-CoV-2 spike pseudotyped lentiviral particles (enough to generate >1,000- fold signal over background, approximately 3 to 4×10^5^ relative light units [RLU]/well) in 96-well plates at a 1:1 ratio (120 µL final volume) and incubated at 37 °C, 5% CO_2_ for 1 hour. The serially diluted serum with SARS-CoV-2 spike pseudotyped lentiviruses was then added to HEK293/ACE2 cells and incubated at 37 °C, 5% CO_2_ for 72 hours. Viral entry was quantified by removing the cell culture supernatant and adding a 1:1 mixture of fresh cell culture media (40 µL) and luciferin reagent (Steady-Luc Firefly Assay Kit, Biotium; 40 µL) to each well. Plates were incubated at RT with gentle shaking (300 rpm) for 5 minutes and luminescence measured using a plate reader (VICTOR Nivo, PerkinElmer). Percent inhibition and 50% neutralising antibody titres (NT50) were calculated as previously described^56^.

## Statistical analysis

All graphs and statistical analyses were undertaken using the GraphPad Prism software (version 10.3.0). Data are represented as mean ± SD. Value of *n* represents the number of animals used and the number of independent experiments conducted, as indicated in the figure legends. Statistical analysis of P < 0.05 was considered significant, determined by unpaired t-test or one-way analysis of variance (ANOVA) with Tukey’s multiple-comparison test, as specified in the relevant figure legends. Schematics in figures were created with BioRender.

## Data availability

All data generated or analysed during this study are included in this published article and the supplementary information files. Any additional information required to reanalyse the reported data is available from the lead contact upon request.

## Author contributions

ALM, TO, WMP and LMC conceived the study and designed experiments. ALM, JFC, KHB, ORP, NCM, IM and LW performed immunogenicity and antigen expression experiments. ALM and JK performed pseudovirus neutralisation assays. SLD and AJF encapsulated vaccines in LNPs and GFP acquired funding for this work. TWB and LW expressed and purified the RBD and NP proteins. ALM, WMP and LMC performed data analysis. WMP and LMC acquired funding, supervised the research and were the project managers. ALM, WMP and LMC wrote the manuscript. All authors read and approved the final manuscript.

## Acknowledgements

ALM is the grateful recipient of a Wellington Doctoral Scholarship. ALM’s doctoral work is also supported by the Rex & Betty Coker Foundation, the Margaret Ann Tibbles Charitable Trust, and the AJ and JC Heine Charitable Trust. We thank the Malaghan Institute of Medical Research’s Biomedical Research Unit for animal care and its Hugh Green Technology Centre for technical assistance with flow cytometry. We thank Dr Rebecca McKenzie and the RNA Technology Platform RNA production team for advice and support on RNA production and quality control. This study was funded by the Maurice Wilkins Centre [grant number MWC4032] and the New Zealand Ministry of Business, Innovation and Employment’s RNA Technology Platform [grant number E4660]. The funders played no role in study design, data collection, analysis and interpretation of data, or the writing of this manuscript.

## Competing interests

All authors declare no financial or non-financial competing interests.

## Notes

### Competing Interest Statement

The authors have declared no competing interest.

## References

1. Wesselhoeft, R. A. et al. RNA Circularization Diminishes Immunogenicity and Can Extend Translation Duration *In Vivo*. Mol. Cell 74, 508–520.e4 (2019).

2. Qu, L. et al. Circular RNA vaccines against SARS-CoV-2 and emerging variants. Cell 185, 1728–1744.e16 (2022).

3. Wesselhoeft, R. A., Kowalski, P. S. & Anderson, D. G. Engineering circular RNA for potent and stable translation in eukaryotic cells. Nat. Commun. 9, 2629 (2018).

4. Hornung, V. et al. 5′-Triphosphate RNA Is the Ligand for RIG-I. Science 314, 994–997 (2006).

5. Rehwinkel, J. & Gack, M. U. RIG-I-like receptors: their regulation and roles in RNA sensing. Nat. Rev. Immunol. 20, 537–551 (2020).

6. Lu, C. et al. The Structural Basis of 5′ Triphosphate Double-Stranded RNA Recognition by RIG-I C-Terminal Domain. Structure 18, 1032–1043 (2010).

7. Jiang, F. et al. Structural basis of RNA recognition and activation by innate immune receptor RIG-I. Nature 479, 423–427 (2011).

8. C Chen, Y. G., et al. Sensing Self and Foreign Circular RNAs by Intron Identity. Mol. Cell 67, 228–238.e5 (2017).

9. Zhang, Y. et al. Rational Design and Immunological Mechanisms of Circular RNA-Based Vaccines: Emerging Frontiers in Combating Pathogen Infection. Vaccines 13, 563 (2025).

10. Baden, L. R. et al. Efficacy and Safety of the mRNA-1273 SARS-CoV-2 Vaccine. N. Engl. J. Med. 384, 403–416 (2021).

11. Corbett, K. S. et al. Evaluation of the mRNA-1273 Vaccine against SARS-CoV-2 in Nonhuman Primates. N. Engl. J. Med. 383, 1544–1555 (2020).

12. P Polack, F. P., et al. Safety and Efficacy of the BNT162b2 mRNA Covid-19 Vaccine. N. Engl. J. Med. 383, 2603–2615 (2020).

13. Chaudhary, N., Weissman, D. & Whitehead, K. A. mRNA vaccines for infectious diseases: principles, delivery and clinical translation. Nat. Rev. Drug Discov. 20, 817–838 (2021).

14. Pardi, N. & Krammer, F. mRNA vaccines for infectious diseases — advances, challenges and opportunities. Nat. Rev. Drug Discov. 23, 838–861 (2024).

15. Yaremenko, A. V., Khan, M. M., Zhen, X., Tang, Y. & Tao, W. Clinical advances of mRNA vaccines for cancer immunotherapy. Med 6, 100562 (2025).

16. Liu, C. et al. mRNA-based cancer therapeutics. Nat. Rev. Cancer 23, 526–543 (2023).

17. Kato, H. et al. Length-dependent recognition of double-stranded ribonucleic acids by retinoic acid–inducible gene-I and melanoma differentiation–associated gene 5. J. Exp. Med. 205, 1601–1610 (2008).

18. Karikó, K., Buckstein, M., Ni, H. & Weissman, D. Suppression of RNA Recognition by Toll-like Receptors: The Impact of Nucleoside Modification and the Evolutionary Origin of RNA. Immunity 23, 165–175 (2005).

19. Karikó, K. et al. Incorporation of Pseudouridine Into mRNA Yields Superior Nonimmunogenic Vector With Increased Translational Capacity and Biological Stability. Mol. Ther. 16, 1833–1840 (2008).

20. Andries, O. et al. N^1^-methylpseudouridine-incorporated mRNA outperforms pseudouridine-incorporated mRNA by providing enhanced protein expression and reduced immunogenicity in mammalian cell lines and mice. J. Control. Release 217, 337–344 (2015).

21. Karikó, K., Muramatsu, H., Ludwig, J. & Weissman, D. Generating the optimal mRNA for therapy: HPLC purification eliminates immune activation and improves translation of nucleoside-modified, protein-encoding mRNA. Nucleic Acids Res. 39, e142–e142 (2011).

22. Puttaraju, M. & Been, M. Group I permuted intron-exon (PIE) sequences self-splice to produce circular exons. Nucleic Acids Res. 20, 5357–5364 (1992).

23. Chen, C. & Sarnow, P. Initiation of Protein Synthesis by the Eukaryotic Translational Apparatus on Circular RNAs. Science 268, 415–417 (1995).

24. Huang, K. et al. Circular mRNA Vaccine against SARS-COV-2 Variants Enabled by Degradable Lipid Nanoparticles. ACS Appl. Mater. Interfaces 17, 4699–4710 (2025).

25. Singh, O. N. et al. Comparison of immunogenicity and protection efficacy of self-amplifying and circular mRNA vaccines against SARS-CoV-2. iScience 28, 113498 (2025).

26. Liu, X. et al. A single-dose circular RNA vaccine prevents Zika virus infection without enhancing dengue severity in mice. Nat. Commun. 15, 8932 (2024).

27. Yue, X. et al. CircRNA based multivalent neuraminidase vaccine induces broad protection against influenza viruses in mice. npj Vaccines 9, 170 (2024).

28. Zhang, W. et al. Bivalent circular RNA vaccines against porcine epidemic diarrhea virus and transmissible gastroenteritis virus. Front. Immunol. 16, 1562865 (2025).

29. Amaya, L., et al. Circular RNA vaccine induces potent T cell responses. Proc. Natl Acad. Sci. USA 120, e2302191120 (2023).

30. Fan, B. et al. Mechanistic insights into circularization via *Anabaena* group I intron-based scarless circular RNA. Mol. Ther. Nucleic Acids 36, 102626 (2025).

31. Chen, L. et al. Development and comprehensive evaluation of scarless circularization systems for circular RNA therapeutics. Mol. Ther. Nucleic Acids 36, 102587 (2025).

32. Qiu, Z. et al. Clean-PIE: a novel strategy for efficiently constructing precise circRNA with thoroughly minimized immunogenicity to direct potent and durable protein expression. Preprint at https://www.biorxiv.org/content/10.1101/2022.06.20.496777v2 (2022).

33. Zhang, Z. et al. Mitigating Cellular Dysfunction by Addressing Contaminants in Synthetic circRNA. Preprint at https://www.biorxiv.org/content/10.1101/2024.09.17.613157v1 (2024).

34. Yang, Y. et al. Extensive translation of circular RNAs driven by *N*^6^-methyladenosine. Cell Res. 27, 626–641 (2017).

35. Chen, Y. G., et al. *N*6-Methyladenosine Modification Controls Circular RNA Immunity. Mol. Cell 76, 96–109.e9 (2019).

36. Nelson, J. et al. Impact of mRNA chemistry and manufacturing process on innate immune activation. Sci. Adv. 6, eaaz6893 (2020).

37. Mu, X., Greenwald, E., Ahmad, S. & Hur, S. An origin of the immunogenicity of *in vitro* transcribed RNA. Nucleic Acids Res. 46, 5239–5249 (2018).

38. Alameh, M.-G. et al. Lipid nanoparticles enhance the efficacy of mRNA and protein subunit vaccines by inducing robust T follicular helper cell and humoral responses. Immunity 54, 2877–2892.e7 (2021).

39. Cheng, F. et al. Study on the Characterization and Degradation Pattern of Circular RNA Vaccines Using an HPLC Method. Chemosensors 12, 120 (2024).

40. Liu, C.-X. et al. RNA circles with minimized immunogenicity as potent PKR inhibitors. Mol. Cell 82, 420–434.e6 (2022).

41. Anderson, B. R. et al. Incorporation of pseudouridine into mRNA enhances translation by diminishing PKR activation. Nucleic Acids Res. 38, 5884–5892 (2010).

42. Wang, Z. et al. Reducing cell intrinsic immunity to mRNA vaccine alters adaptive immune responses in mice. Mol. Ther. Nucleic Acids 34, 102045 (2023).

43. Bernard, M.-C. et al. The impact of nucleoside base modification in mRNA vaccine is influenced by the chemistry of its lipid nanoparticle delivery system. Mol. Ther. Nucleic Acids 32, 794–806 (2023).

44. Danz, H. et al. Synergistic effect of nucleoside modification and ionizable lipid composition on translation and immune responses to mRNA vaccines. npj Vaccines 10, 212 (2025).

45. Wang, X., et al. N^6^-methyladenosine Modulates Messenger RNA Translation Efficiency. Cell 161, 1388–1399 (2015).

46. Zhou, J. et al. Dynamic m^6^A mRNA methylation directs translational control of heat shock response. Nature 526, 591–594 (2015).

47. Meyer, K. D. et al. 5′ UTR m^6^A Promotes Cap-Independent Translation. Cell 163, 999–1010 (2015).

48. Victora, G. D. & Nussenzweig, M. C. Germinal Centers. Annu. Rev. Immunol. 40, 413–442 (2022).

49. Tam, H. H., et al. Sustained antigen availability during germinal center initiation enhances antibody responses to vaccination. Proc. Natl Acad. Sci. USA 113, (2016).

50. Wan, J. et al. Circular RNA vaccines with long-term lymph node-targeting delivery stability after lyophilization induce potent and persistent immune responses. mBio 15, e01775–23 (2024).

51. Wilkins, M. R. et al. Protein Identification and Analysis Tools in the ExPASy Server. Methods Mol. Biol. 112, 531–552 (1999).

52. McKenzie, R. E., Minnell, J. J., Ganley, M., Painter, G. F. & Draper, S. L. mRNA Synthesis and Encapsulation in Ionizable Lipid Nanoparticles. Curr. Protoc. 3, e898 (2023).

53. Frey, A., Di Canzio, J. & Zurakowski, D. A statistically defined endpoint titer determination method for immunoassays. J. Immunol. Methods 221, 35–41 (1998).

54. Montgomerie, I. et al. Incorporation of SARS-CoV-2 spike NTD to RBD protein vaccine improves immunity against viral variants. iScience 26, 106256 (2023).

55. Crawford, K. H. D. et al. Protocol and Reagents for Pseudotyping Lentiviral Particles with SARS-CoV-2 Spike Protein for Neutralization Assays. Viruses 12, 513 (2020).

56. Nie, J. et al. Quantification of SARS-CoV-2 neutralizing antibody by a pseudotyped virus-based assay. Nat. Protoc. 15, 3699–3715 (2020).

