## Supplementary Information for "Circular RNA vaccine performance is determined by RNA quality and epitranscriptomic tuning rather than innate activation"

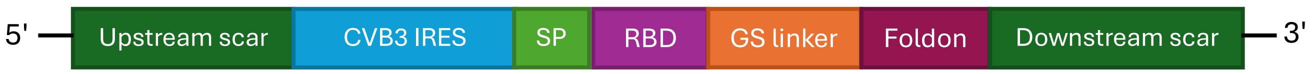


AAAAUCCGUUGACCUUAAACGGUCGUGUGGGUUCAAGUCCCUCCACCCCCACGCCGGAAACG
CAAUAGCCGAAAAACAAAAAACAAAAAAAACAAAAAAAAAACCAAAAAAACAAAACACAUUA
AAACAGCCUGUGGGUUGAUCCCACCCACAGGCCCAUUGGGCGCUAGCACUCUGGUAUCACGG
UACCUUUGUGCGCCUGUUUUAUACCCCCUCCCCCAACUGUAACUUAGAAGUAACACACACCG
AUCAACAGUCAGCGUGGCACACCAGCCACGUUUUGAUCAAGCACUUCUGUUACCCCGGACUG
AGUAUCAAUAGACUGCUCACGCGGUUGAAGGAGAAAGCGUUCGUUAUCCGGCCAACUACUUC
GAAAAACCUAGUAACACCGUGGAAGUUGCAGAGUGUUUCGCUCAGCACUACCCCAGUGUAGA
UCAGGUCGAUGAGUCACCGCAUUCCCCACGGGCGACCGUGGCGGUGGCUGCGUUGGCGGCCU
GCCCAUGGGGAAACCCAUGGGACGCUCUAAUACAGACAUGGUGCGAAGAGUCUAUUGAGCUA
GUUGGUAGUCCUCCGGCCCCUGAAUGCGGCUAAUCCUAACUGCGGAGCACACACCCUCAAGC
CAGAGGGCAGUGUGUCGUAACGGGCAACUCUGCAGCGGAACCGACUACUUUGGGUGUCCGUG
UUUCAUUUUAUUCCUAUACUGGCUGCUUAUGGUGACAAUUGAGAGAUCGUUACCAUAUAGCU
AUUGGAUUGGCCAUCCGGUGACUAAUAGAGCUAUUAUAUCCCUUUGUUGGGUUUAUACCA
CUUAGCUUGAAAGAGGUUAAAACAUUACAAUUCAUUGUUAAGUUGAAUACAGCAAA**AUG**GAC
GCUAUGAAGAGGGGACUGUGCUGCGUGCUGCUGCUGUGCGGAGCCGUGUUCGUGAGCCCAUC
CCAGGAGAUCCACGCCAGGUUCAGGAGGAGGGUGCAGCCUACCGAGUCUAUCGUGCGGUUCC
CUAACAUCACAAACCUGUGCCCAUUCGGCGAGGUGUUUAACGCCACCAGGUUUGCUAGCGUG
UACGCCUGGAACCGCAAGAGGAUCUCUAACUGCGUGGCUGACUACAGCGUGCUGUACAACAG
CGCCAGCUUCAGCACCUUCAAGUGCUACGGCGUGAGCCCCACCAAGCUGAACGACCUGUGCU
UCACAAACGUGUACGCUGACUCCUUUGUGAUCCGCGGAGAUGAGGUGAGGCAGAUCGCUCCA
GGCCAGACCGGAAAGAUCGCCGACUACAACUACAAGCUGCCCGACGAUUUCACAGGCUGCGU
GAUCGCCUGGAACUCCAACAACCUGGAUUCUAAAGUGGGCGGAAACUACAACUACAGAUACC
GGCUGUUCCGCAAGUCCAACCUGAAGCCUUUUGAGAGGGAUAUCUCCACCGAGAUCUACCAG
GCCGGCUCUAAGCCAUGCAACGGCGUGGAGGGAUUCAACUGUUACUUUCCCCUGCAGUCUUA
CGGAUUCCAGCCUACAAACGGCGUGGGAUACCAGCCAUACAGAGUGGUGGUGCUGUCUUUUG
AGCUGCUGCACGCUCCAGCUACCGUGUGCGGACCCAAGAAGUCCACAAACCUGGUGAAGAAC
AAGUGCGUGAACUUCGGCGGAGGAGGCAGCGGAGGAGGAGGAUCUGGCAGCGGAUACAUCCC
CGAGGCUCCUAGAGACGGCCAGGCCUACGUGCGGAAGGAUGGAGAGUGGGUGCUGCUGAGCA
CCUUUCUGGGACGGUCC**UAA**AAAAAACAAAAAACAAAACGGCUAUUAUGCGUUACCGGCGAG
ACGCUACGGACUU

**Supplementary Figure 1.** CircRNA$^{RBD}$ constructs (unmodified; 1% and 5% m$^6$A randomly incorporated during in vitro transcription). Colour coding in the RNA sequence corresponds to the diagram at the top of the figure. Upstream and downstream scars (dark green) are the intronic sequences that remain after circularisation. The internal ribosome entry site (IRES) was from coxsackie virus B3 (CVB3) and mediates cap-independent translation. The signal peptide (SP) is from tissue plasminogen (tPA) and ensures secretion of the encoded receptor binding domain (RBD) antigen. The GS linker sequence encodes a glycine-serine linker. To improve immunogenicity and stability of the RBD antigen, the trimerization motif of bacteriophage T4 fibritin protein (Foldon) was incorporated at the C-terminus. Bolded letters indicate start or stop codons for the encoded antigen (AUG and UAA, respectively).

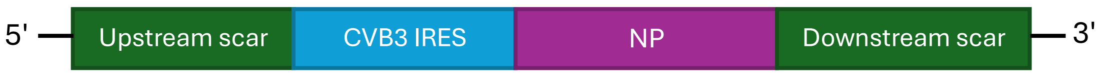


AAAAUCCGUUGACCUUAAACGGUCGUGUGGGUUCAAGUCCCUCCACCCCCACGCCGGAAACG
CAAUAGCCGAAAAACAAAAAACAAAAAAAACAAAAAAAAAACCAAAAAAACAAAACACAUUA
AAACAGCCUGUGGGUUGAUCCCACCCACAGGCCCAUUGGGCGCUAGCACUCUGGUAUCACGG
UACCUUUGUGCGCCUGUUUUAUACCCCCUCCCCCAACUGUAACUUAGAAGUAACACACACCG
AUCAACAGUCAGCGUGGCACACCAGCCACGUUUUGAUCAAGCACUUCUGUUACCCCGGACUG
AGUAUCAAUAGACUGCUCACGCGGUUGAAGGAGAAAGCGUUCGUUAUCCGGCCAACUACUUC
GAAAAACCUAGUAACACCGUGGAAGUUGCAGAGUGUUUCGCUCAGCACUACCCCAGUGUAGA
UCAGGUCGAUGAGUCACCGCAUUCCCCACGGGCGACCGUGGCGGUGGCUGCGUUGGCGGCCU
GCCCAUGGGGAAACCCAUGGGACGCUCUAAUACAGACAUGGUGCGAAGAGUCUAUUGAGCUA
GUUGGUAGUCCUCCGGCCCCUGAAUGCGGCUAAUCCUAACUGCGGAGCACACACCCUCAAGC
CAGAGGGCAGUGUGUCGUAACGGGCAACUCUGCAGCGGAACCGACUACUUUGGGUGUCCGUG
UUUCAUUUUAUUCCUAUACUGGCUGCUUAUGGUGACAAUUGAGAGAUCGUUACCAUAUAGCU
AUUGGAUUGGCCAUCCGGUGACUAAUAGAGCUAUUAUAUAUCCCUUUGUUGGGUUUAUACCA
CUUAGCUUGAAAGAGGUUAAAACAUUACAAUUCAUUGUUAAGUUGAAUACAGCAAA**AUG**GCC
UCCCAGGGCACCAAGCGGUCUUACGAGCAGAUGGAGACCGACGGAGAGAGACAGAACGCCAC
AGAGAUCCGGGCUUCCGUGGGCAAGAUGAUCGGCGGAAUCGGAAGAUUCUACAUCCAGAUGU
GCACCGAGCUGAAGCUGUCUGAUUACGAGGGCAGACUGAUCCAGAACAGCCUGACAAUCGAG
CGGAUGGUGCUGUCCGCCUUUGACGAGAGGAGAAACAAGUACCUGGAGGAGCACCCUUCCGC
UGGCAAGGACCCCAAGAAGACCGGAGGACCAAUCUACAGGAGGGUGAACGGAAAGUGGAUGC
GGGAGCUGAUCCUGUACGACAAGGAGGAGAUCAGGAGAAUCUGGCGCCAGGCCAACAACGGC
GACGAUGCCACCGCUGGACUGACACACAUGAUGAUCUGGCACUCUAACCUGAACGACGCCAC
CUACCAGAGGACAAGAGCUCUGGUGAGAACCGGCAUGGACCCCCGGAUGUGCUCCCUGAUGC
AGGGAUCUACACUGCCACGGCGCAGCGGAGCUGCUGGAGCUGCUGUGAAGGGCGUGGGAACC
AUGGUCAUGGAGCUGGUGCGGAUGAUCAAGCGCGGCAUCAACGACAGGAACUUCUGGAGAGG
CGAGAACGGACGGAAGACCCGCAUCGCCUACGAGAGGAUGUGCAACAUCCUGAAGGGCAAGU
UUCAGACAGCCGCUCAGAAGGCCAUGAUGGACCAGGUGAGGGAGUCUAGAGAUCCCGGCAAC
GCUGAGUUCGAGGAUCUGACAUUUCUGGCCAGGAGCGCCCUGAUCCUGAGGGGAAGCGUGGC
UCACAAGUCCUGCCUGCCUGCUUGCGUGUACGGACCAGCUGUGGCUAGCGGAUACGACUUCG
AGCGCGAGGGCUACUCCCUGGUGGGAAUCGAUCCUUUUAGGCUGCUGCAGAACUCUCAGGUG
UACAGCCUGAUCAGACCAAACGAGAACCCCGCCCACAAGUCUCAGCUGGUGUGGAUGGCUUG
UCACAGCGCCGCUUUCGAGGACCUGAGGGUGCUGUCCUUUAUCAAGGGCACCAAGGUGGUGC
CAAGAGGCAAGCUGUCCACACGGGGAGUGCAGAUCGCCUCUAACGAGAACAUGGAGACCAUG
GAGAGCUCCACACUGGAGCUGCGGUCCCGCUACUGGGCUAUCAGGACCAGAUCUGGCGGAAA
CACAAACCAGCAGAGAGCCUCUGCUGGCCAGAUCAGCAUCCAGCCCACCUUCAGCGUGCAGC
GCAACCUGCCUUUUGAUAGGACCACAGUGAUGGCCGCUUUCACCGGCAACACAGAGGGACGG
ACCUCCGACAUGCGCACAGAGAUCAUCAGGAUGAUGGAGUCUGCCAGGCCAGAGGAUGUGAG
CUUCCAGGGAAGGGGAGUGUUUGAGCUGAGCGACGAGAAGGCCGCUUCCCCCAUCGUGCCUU
CCUUCGAUAUGUCUAACGAGGGCAGCUACUUCUUUGGAGACAACGCCGAGGAGUACGAUAAC
AGCCACCACCACCACCACCAC**UAA**AAAAAACAAAAAACAAAACGGCUAUUAUGCGUUACCGG
CGAGACGCUACGGACUU

**Supplementary Figure 2:** CircRNA$^{NP}$ constructs (unmodified; 1% m$^6$A randomly incorporated during in vitro transcription). Colour coding in the RNA sequence corresponds to the diagram at the top of the figure. Upstream and downstream scars (dark green) are the intronic sequences that remain after circularisation. The internal ribosome entry site (IRES) was from coxsackie virus B3 (CVB3) and mediates cap-independent translation. Bolded letters indicate start or stop codons for the encoded antigen (AUG and UAA, respectively).

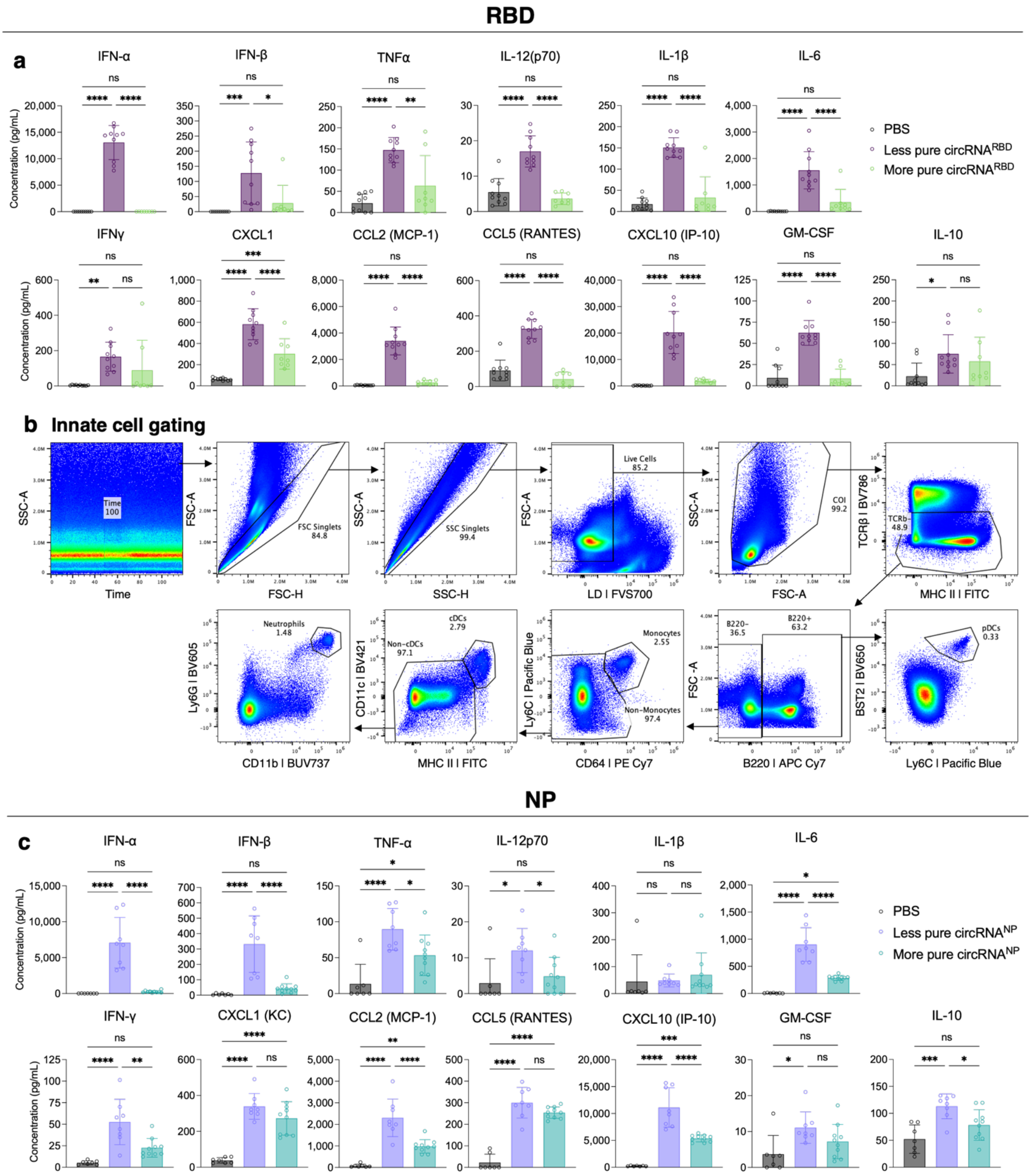


**Supplementary Figure 3: CircRNA is reactogenic when contaminants are present.** (**a**) Serum cytokine responses taken at six hours post-immunisation with circRNA$^{RBD}$ vaccines. (**b**) Representative gating strategy to define innate cell subsets. (**c**) Serum cytokine responses taken at six hours post-immunisation with circRNA$^{NP}$ vaccines. For cytokine data in (**a** and **c**), top row are raw data of the cytokines depicted in Figure 1c and f. Bottom row are other chemokines and cytokines involved in the anti-viral response. Data are a combination of $n$=2 experiments with $n$=4-5 mice per group, per experiment. Graphs are depicted as mean ± SD. One-way ANOVA with Tukey's multiple comparisons test was performed, with ns = not significant; * P < 0.05; ** P < 0.01; *** P < 0.001; **** P < 0.0001.

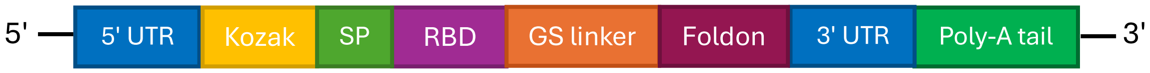


GGGAAAUAAGAGAGAAAAGAAGAGUAAGAAAUAUAAGACCCCGGCGCCGCCACCGCCAC
C**AUG**GACGCUAUGAAGAGGGGACUGUGCUGCGUGCUGCUGCUGUGCGGAGCCGUGUUCGUGA
GCCCAUCCCAGGAGAUCCACGCCAGGUUCAGGAGGAGGGUGCAGCCUACCGAGUCUAUCGUG
CGGUUCCCUAACAUCACAAACCUGUGCCCAUUCGGCGAGGUGUUUAACGCCACCAGGUUUGC
UAGCGUGUACGCCUGGAACCGCAAGAGGAUCUCUAACUGCGUGGCUGACUACAGCGUGCUGU
ACAACAGCGCCAGCUUCAGCACCUUCAAGUGCUACGGCGUGAGCCCCACCAAGCUGAACGAC
CUGUGCUUCACAAACGUGUACGCUGACUCCUUUGUGAUCCGCGGAGAUGAGGUGAGGCAGAU
CGCUCCAGGCCAGACCGGAAAGAUCGCCGACUACAACUACAAGCUGCCCGACGAUUUCACAG
GCUGCGUGAUCGCCUGGAACUCCAACAACCUGGAUUCUAAAGUGGGCGGAAACUACAACUAC
AGAUACCGGCUGUUCCGCAAGUCCAACCUGAAGCCUUUUGAGAGGGAUAUCUCCACCGAGAU
CUACCAGGCCGGCUCUAAGCCAUGCAACGGCGUGGAGGGAUUCAACUGUUACUUUCCCCUGC
AGUCUUACGGAUUCCAGCCUACAAACGGCGUGGGAUACCAGCCAUACAGAGUGGUGGUGCUG
UCUUUUGAGCUGCUGCACGCUCCAGCUACCGUGUGCGGACCCAAGAAGUCCACAAACCUGGU
GAAGAACAAGUGCGUGAACUUCGGCGGAGGAGGCAGCGGAGGAGGAGGAUCUGGCAGCGGAU
ACAUCCCCGAGGCUCCUAGAGACGGCCAGGCCUACGUGCGGAAGGAUGGAGAGUGGGUGCUG
CUGAGCACCUUUCUGGGACGGUCC**UAA**UAGGCUGGAGCCUCGGUGGCCUAGCUUCUUGCCCC
UUGGGCCUCCCCCCAGCCCCUCCUCCCCUUCCUGCACCCGUACCCCCGUGGUCUUUGAAUAA
AGUCUGAGUGGGCGGCAACUAGUAAAAAAAAAAAAAAAAAAAAAAAAAAAAAAAAAAAA
AAAAAAAAAAAAAAAAAAAAAAAAAAAAAAAAAAAAAAAAAAAAAAAAAAAAAAAAAA

**Supplementary Figure 4:** Sequence of m1Ψ-mRNA$^{RBD}$. mRNA constructs incorporated a Cap 1 structure (m$^7$GpppNmN) at the 5' end to facilitate cap-dependent translation. Colour coding in the RNA sequence corresponds to the diagram at the top of the figure. Untranslated regions (UTR) were incorporated at both the 5' and 3' ends. The signal peptide (SP) is from tissue plasminogen (tPA) and ensures secretion of the encoded receptor binding domain (RBD) antigen. The GS linker sequence encodes a glycine-serine linker. To improve immunogenicity and stability of the RBD antigen, the trimerization motif of bacteriophage T4 fibritin protein (Foldon) was incorporated at the C-terminus. The polyA tail (100A) was encoded in the plasmid backbone. Bolded letters indicate start or stop codons (AUG and UAA, respectively). During mRNA synthesis, *N*$^1$-methylpseudouridine (m1Ψ) was incorporated instead of uridine.

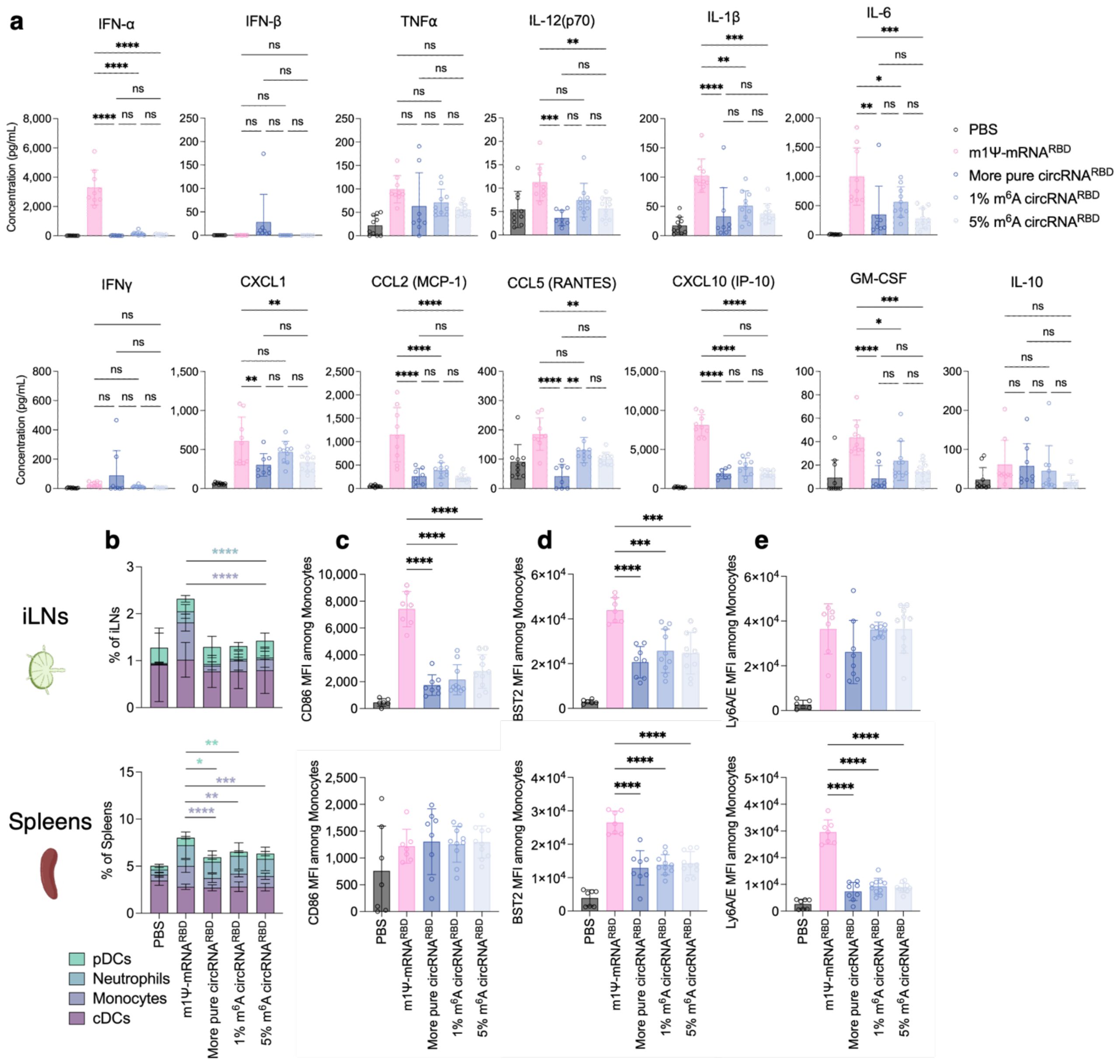


**Supplementary Figure 5: Reactogenicity of m$^6$A-modified circRNA$^{RBD}$.** (**a**) Serum cytokine responses taken at six hours post-immunisation. Top row are raw data of the cytokines depicted in Figure 2b. Bottom row are other chemokines and cytokines involved in the anti-viral response. (**b**) Stacked bar graphs highlighting innate cell frequencies in iLNs (top) and spleens (bottom). MFI of (**c**) CD86, (**d**) BST2, and (**e**) Ly6A/E on monocytes in iLNs (top) and spleens (bottom). Data are a combination of $n$=2-4 experiments with $n$=3-5 mice per group, per experiment. Graphs are depicted as mean ± SD. One-way ANOVA with Tukey's multiple comparisons test was performed, with ns = not significant; * $P < 0.05$; ** $P < 0.01$; *** $P < 0.001$; **** $P < 0.0001$.

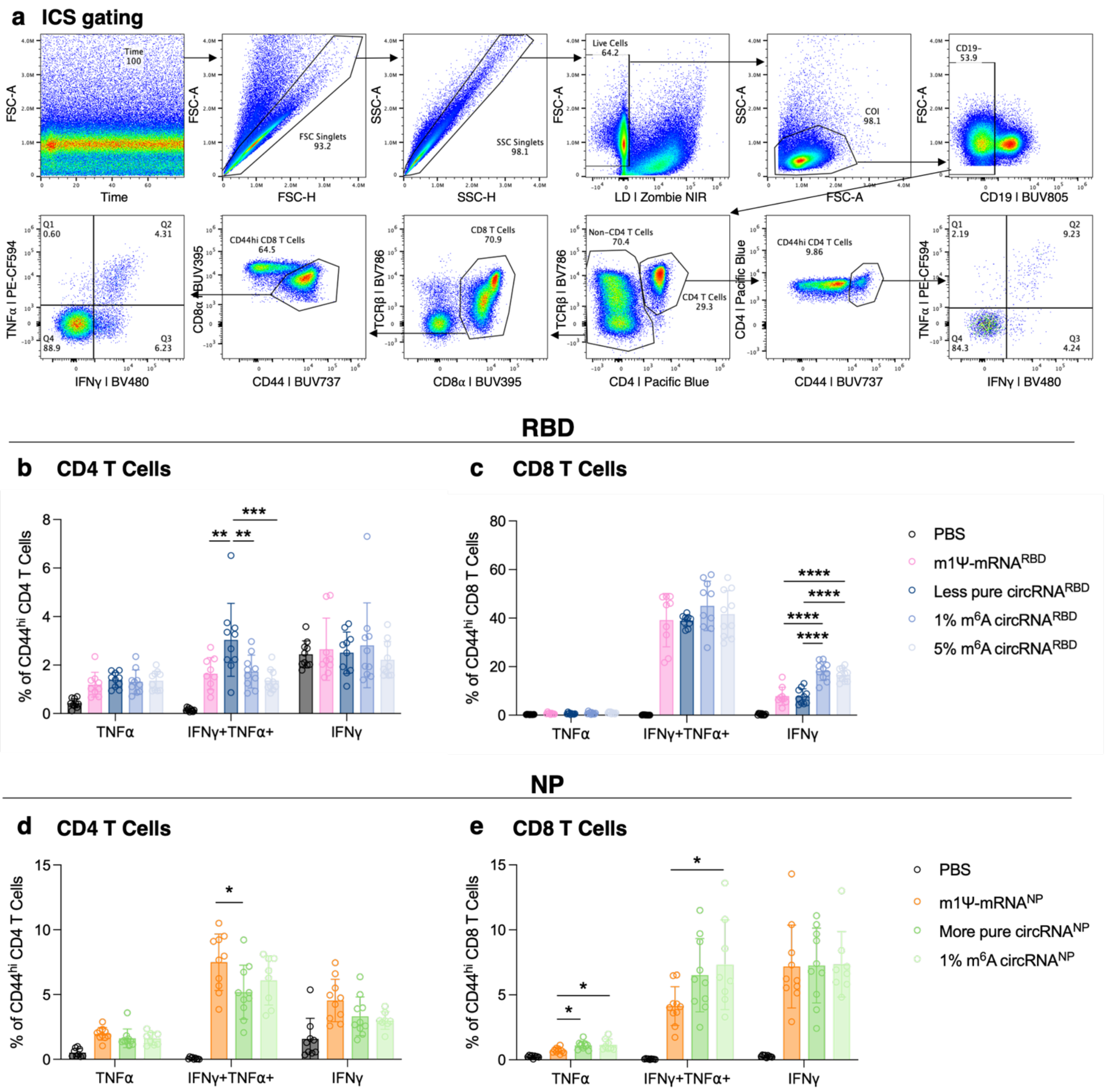


**Supplementary Figure 6: CircRNA and linear mRNA elicit comparable adaptive immune magnitude independent of innate immune reactogenicity.** (**a**) Representative gating strategy to define cytokine-producing CD4 and CD8 T cell subsets. Frequency of cytokine-producing (**b**) CD44$^{hi}$ CD4 T cells and (**c**) CD44$^{hi}$ CD8 T cells following S1 peptide stimulation of spleens from RBD-immunised mice. Frequency of cytokine-producing (**d**) CD44$^{hi}$ CD4 T cells and (**e**) CD44$^{hi}$ CD8 T cells following NP peptide stimulation of spleens from NP-immunised mice. Data are a combination of *n*=2 experiments with *n*=4-5 mice per group, per experiment. Graphs are depicted as mean ± SD. One-way ANOVA with Tukey's multiple comparisons test was performed, with * P < 0.05; ** P < 0.01; *** P < 0.001; **** P < 0.0001.

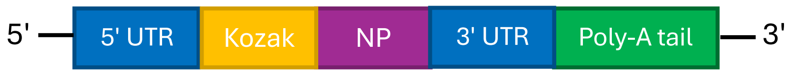


GGGAAAUAAGAGAGAAAAGAAGAGUAAGAAGAAAUAUAAGACCCCGGCGCCGCCACCGCCAC
C**AUG**GCGUCCCAAGGCACCAAACGGUCUUACGAACAGAUGGAGACUGAUGGAGAACGCCAGA
AUGCCACUGAAAUCAGAGCAUCCGUCGGAAAAAUGAUUGGUGGAAUUGGACGAUUCUACAUC
CAAAUGUGCACAGAACUUAAACUCAGUGAUUAUGAGGGACGGUUGAUCCAAAACAGCUUAAC
AAUAGAGAGAAUGGUGCUCUCUGCUUUUGACGAAAGGAGAAAUAAAUACCUGGAAGAACAUC
CCAGUGCGGGGAAGGACCCUAAGAAAACUGGAGGACCUAUAUACAGAAGAGUAAACGGAAAG
UGGAUGAGAGAACUCAUCCUUUAUGACAAAGAAGAAAUAAGGCGAAUCUGGCGCCAAGCUAA
UAAUGGUGACGAUGCAACGGCUGGUCUGACUCACAUGAUGAUCUGGCAUUCCAAUUUGAAUG
AUGCAACUUAUCAGAGGACAAGGGCUCUUGUUCGCACCGGAAUGGACCCCAGGAUGUGCUCU
CUGAUGCAAGGUUCAACUCUCCCUAGGAGGUCUGGAGCCGCAGGUGCUGCAGUCAAAGGAGU
UGGAACAAUGGUGAUGGAAUUGGUCAGGAUGAUCAAACGUGGGAUCAAUGAUCGGAACUUCU
GGAGGGGUGAGAAUGGACGAAAAACAAGAAUUGCUUAUGAAAGAAUGUGCAACAUUCUCAAA
GGGAAAUUUCAAACUGCUGCACAAAAAGCAAUGAUGGAUCAAGUGAGAGAGAGCCGGGACCC
AGGGAAUGCUGAGUUCGAAGAUCUCACUUUUCUAGCACGGUCUGCACUCAUAUUGAGAGGGU
CGGUUGCUCACAAGUCCUGCCUGCCUGCCUGUGUGUAUGGACCUGCCGUAGCCAGUGGGUAC
GACUUUGAAAGAGAGGGAUACUCUCUAGUCGGAAUAGACCCUUUCAGACUGCUUCAAAACAG
CCAAGUGUACAGCCUAAUCAGACCAAAUGAGAAUCCAGCACACAAGAGUCAACUGGUGUGGA
UGGCAUGCCAUUCUGCCGCAUUUGAAGAUCUAAGAGUAUUGAGCUUCAUCAAAGGGACGAAG
GUGGUCCCAAGAGGGAAGCUUUCCACUAGAGGAGUUCAAAUUGCUUCCAAUGAAAAUAUGGA
GACUAUGGAAUCAAGUACACUUGAACUGAGAAGCAGGUACUGGGCCAUAAGGACCAGAAGUG
GAGGAAACACCAAUCAACAGAGGGCAUCUGCGGGCCAAAUCAGCAUACAACCUACGUUCUCA
GUACAGAGAAAUCUCCCUUUUGACAGAACAACCGUUAUGGCAGCAUUCACUGGGAAUACAGA
GGGGAGAACAUCUGACAUGAGGACCGAAAUCAUAAGGAUGAUGGAAAGUGCAAGACCAGAAG
AUGUGUCUUUCCAGGGGCGGGGAGUCUUCGAGCUCUCGGACGAAAAGGCAGCGAGCCCGAUC
GUGCCUUCCUUUGACAUGAGUAAUGAAGGAUCUUAUUUCUUCGGAGACAAUGCAGAGGAGUA
CGACAAUAGCCAUCAUCAUCAUCAUCAC**UAA**UAGGCUGGAGCCUCGGUGGCCUAGCUUCUUG
CCCCUUGGGCCUCCCCCCAGCCCCUCCUCCCCUUCCUGCACCCGUACCCCCGUGGUCUUUGA
AUAAAGUCUGAGUGGGCGGCAUCUAGAAAAAAAAAAAAAAAAAAAAAAAAAAAAAAAAAAAA
AAAAAAAAAAAAAAAAAAAAAAAAAAAAAAAAAAAAAAAAAAAAA

**Supplementary Figure 7:** Sequence of m1Ψ-mRNA$^{NP}$. mRNA constructs incorporated a Cap 1 structure ($m^7$GpppNmN) at the 5' end to facilitate cap-dependent translation. Colour coding in the RNA sequence corresponds to the diagram at the top of the figure. Untranslated regions (UTR) were incorporated at both the 5' and 3' ends. The encoded antigen is the H1N1 influenza nucleoprotein (NP). The polyA tail (100A) was encoded in the plasmid backbone. Bolded letters indicate start or stop codons (AUG and UAA, respectively). During mRNA synthesis, $N^1$-methylpseudouridine (m1Ψ) was incorporated instead of uridine.

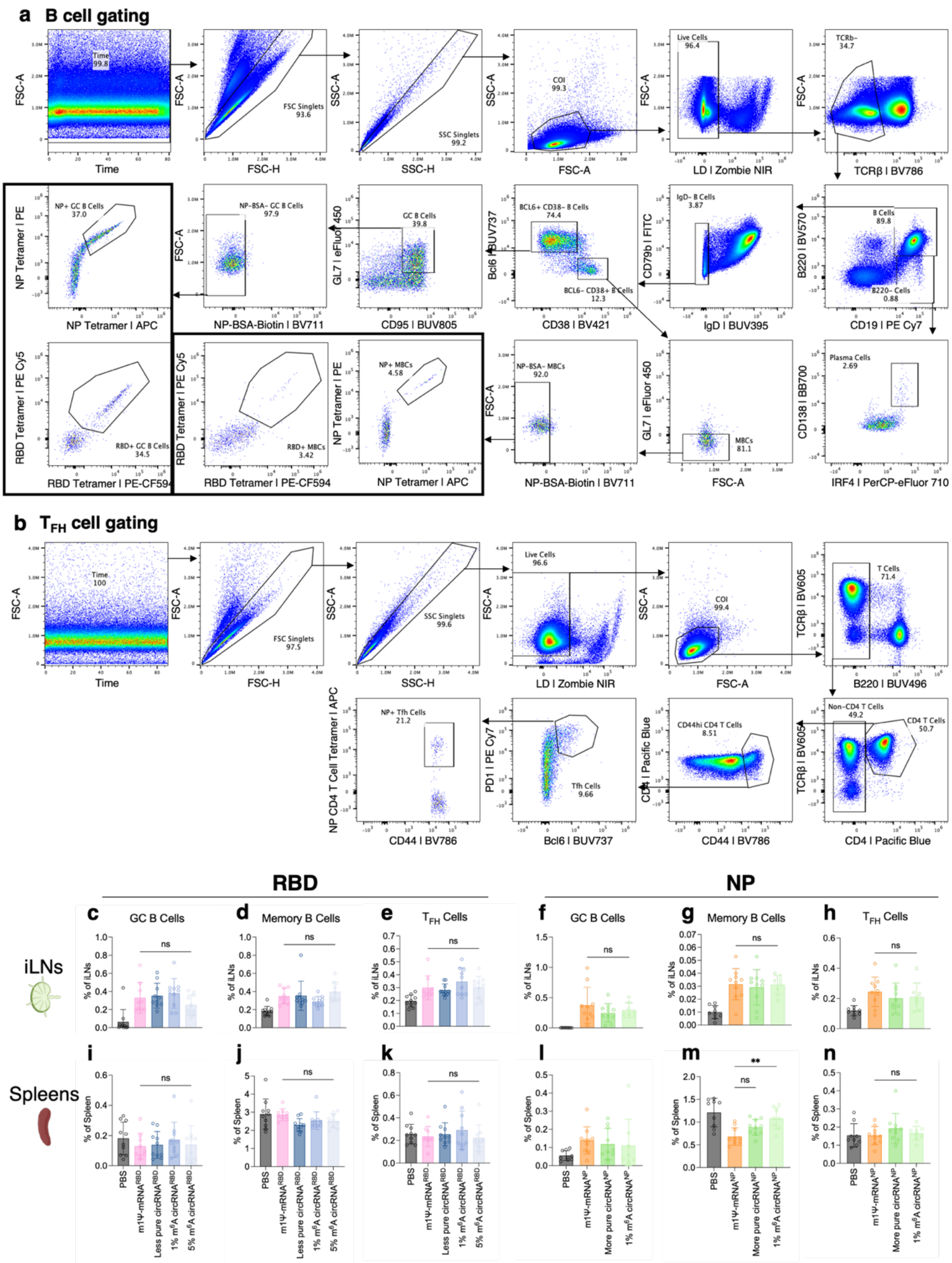


**Supplementary Figure 8: CircRNA vaccination alters the anatomical distribution of germinal centre B cell responses.** (**a**) Representative gating strategy to define antigen-specific B cell populations. (**b**) Representative gating strategy to define $T_{FH}$ cells. Frequency of (**c**) GC B cells, (**d**) memory B cells, and (**e**) $T_{FH}$ cells in iLNs of RBD-immunised mice. Frequency of (**f**) GC B cells, (**g**) memory B cells, and (**h**) $T_{FH}$ cells in iLNs of NP-immunised mice. (**i-n**) Same as (**c-h**) but in spleens. Data are a combination of $n$=2 experiments with $n$=4-5 mice per group, per experiment. Graphs are depicted as mean ± SD. One-way ANOVA with Tukey's multiple comparisons test was performed, with ns = not significant; ** P < 0.01.

**Supplementary Table 1: Relative concentrations of different species in the RNA preparations used in this study.**

| RNA preparation | Correct size[a] | Low MW species[b] | High MW species[c] |
|---|---|---|---|
| **SARS-CoV-2 RBD** | | | |
| circRNA (less pure) | 90.9 % | 5.8 % | 3.4 % |
| circRNA (more pure) | 96.2 % | 0.0 % | 3.8 % |
| circRNA with 1% $m^6A$ | 98.9 % | 0.0 % | 1.1 % |
| circRNA with 5% $m^6A$ | 98.9 % | 0.0 % | 1.1 % |
| mRNA with m1Ψ | 97.7 % | 1.6 % | 0.7 % |
| **Influenza NP** | | | |
| circRNA (less pure) | 87.8 % | 10.6 % | 1.6 % |
| circRNA (more pure) | 98.3 % | 0.0 % | 1.7 % |
| circRNA with 1% $m^6A$ | 92.2 % | 0.0 % | 7.8 % |
| mRNA with m1Ψ | 92.3 % | 5.0 % | 2.6 % |

[a] RNA with the expected size, as determined by the commercial supplier using capillary electrophoresis.

[b] Sum of all RNA species with lower molecular weight (MW) than the desired product.

[c] Sum of all RNA species with higher MW than the desired product.
